# Investigating the apical notch meristem, apical dominance and meristem regeneration in Marchantia polymorpha

**DOI:** 10.1101/2024.02.04.575544

**Authors:** Alan O. Marron

**Affiliations:** Department of Plant Sciences, University of Cambridge, Downing Street, Cambridge CB2 3EA, United Kingdom; Present Address: Oxford Brookes Centre for BioImaging, Oxford Brookes University, Headington Campus, Oxford OX3 0BP, United Kingdom

## Abstract

Meristems are the growth centres of plants, and fundamental in understanding plant development, morphogenesis and vegetative propagation. Across all plant groups, the phytohormone auxin controls meristem maintenance, represses the emergence of new meristems (apical dominance) and mediates cellular reprogramming when new meristems regenerate following removal of existing meristems. The liverwort Marchantia produces clonal propagules (gemmae) featuring two apical notches that develop into functional meristems. This presents a tractable experimental system to study meristem developmental biology. I used laser ablation microscopy to precisely disrupt cells in and around the developing pre-meristem in the apical notches of germinating gemma, finding that the first cell row is indispensable. Within this layer, a contiguous quorum of stem cells is required for activity. Apical notches reorientate in response to damage, demonstrating that the apical notch stem cells act as a communicating population. Feedback from the stem cell population is necessary to maintain notch activity and generate the notch apex. These experiments show communication between notches and regenerating meristems. The apical dominance signal represses cell division and requires both sources and sinks, features of auxin-mediated communication. Central regions of the gemma could transmit these apical dominance signals, but the tissues of the gemma periphery could not. I present a model of Marchantia gemma and apical notch organization, involving intra-, inter-and extra-notch communication. This provides a framework for further study of meristem formation, communication and maintenance in Marchantia and improving knowledge of plant meristems more generally.

**Significance Statement:** Meristems are the growth centres of plants. Understanding how meristems function, how existing meristems stop new meristems emerging (apical dominance) and how cellular reprogramming regenerates meristems is central to understanding plant development. The model plant Marchantia offers a streamlined system whose evolutionary position makes its biology relevant to all plants. Marchantia apical notches display proliferative, regenerative and dominance behaviour typical of meristems. I used laser ablation to precisely disrupt cells in and around the developing pre-meristematic apical notch of the germinating Marchantia gemma, producing an improved model of Marchantia apical notch organization involving intra-and inter-notch communication characteristic of the phytohormone auxin. This will inform future Marchantia research and provides a simplified experimental framework to study more complex meristematic processes.

## Introduction

Cell division in plants occurs in meristematic regions. Meristems contain pluripotent stem cells which continually divide asymmetrically to produce a daughter and a meristematic cell. The cells at the centre of the meristem divide slowly, while those at the periphery divide faster. This moves daughter cells outwards as they mature and differentiate, while meristematic cells remain in the centre, replenishing the stem cell pool. This spatial distinction between division and differentiation allows plant meristems to sustain growth and new organ formation simultaneously (1). It means that meristem size and spacing must be controlled for normal morphogenesis. All plant groups do this by means of positive and negative feedback loops involving signalling molecules such as CLE peptides and phytohormones. Disrupting the biosynthesis or transport of these signalling molecules leads to oversized meristems, incorrectly spaced meristems, or loss of meristem function (2, 3).

Perhaps the most important of these signalling molecules is the phytohormone auxin. In vascular plants, auxin is produced in the shoot apical meristem (SAM) and flows outwards by polar transport, where it is involved in controlling cell expansion and differentiation (4, 5).

Secondary meristem formation is induced by the formation of local auxin maxima and minima arising out of auxin gradients extending from the apical meristems (5). Apically derived auxin inhibits growth of secondary meristems in the buds: this is apical dominance (3). If the SAM is damaged or destroyed, the flow of auxin is disrupted, apical dominance is lifted and these meristems are released from dormancy and begin growing (6). Plants have the capacity to regenerate meristems in response to localized wounding, which changes the auxin:cytokinin ratio around the damage site. In vascular plants, this causes cells from certain tissue types (e.g., embryonic tissue, pericycle) to lose their previous identity, respecify their cell fate and become meristematic tissue (5, 7). This cellular reprogramming to acquire meristem identity involves changes in gene expression, e.g., activation of WIND, WUS, WOX and PLT transcription factors (7). Bryophytes, such as the liverwort *Marchantia polymorpha*, have massive capacity for regeneration: all parts of the plant, and even single cells, can regenerate meristems *de novo* that eventually produce full plants (8–13).

Marchantia grows as a dominant haploid gametophyte thallus, from which develops a diploid sporophyte. Meristems are present in both generations, with some genetic pathways common to both, but other pathways specific to one generation or the other (14–16). The sporophyte lacks an apical meristem, instead growing by cell division along its length (17). The gametophyte thallus has apical meristems located in concave apical notches (Fig. 1*A*), which are the loci of cell proliferation. The apical notch consists of a central apical cell with four cutting faces, surrounded by dorsal and ventral merophytes, plus lateral subapical cells (18, 19). Around each apical notch is a region of cell differentiation. As cells move outwards away from the meristem, division rates reduce, and cell expansion increases. The apical meristems appear early in sporeling development, and their emergence marks the transition from an undifferentiated callus-like protonema into a prothallus, which goes on to form the thallus (9, 20). As the thallus grows the apical notch meristems enlarge and bifurcate, thus producing new thallus branches by dichotomous branching (21). The thallus also forms dorsal outgrowths called gemma cups, within which grow vegetative propagules called gemmae (22). Gemmae have two apical notches located at either end (Fig. 1*B*). Gemmae remain dormant until removal from the gemma cup, at which point they germinate. Immediately after germination the apical notches are in effect pre-meristems, with dorso-ventral and left-right symmetry. This could be considered equivalent to the developing SAM of the late torpedo stage angiosperm embryo, where a stem-cell containing notch, but not yet a morphologically mature meristem, has formed between two cotyledons (23). Over the course of the next five days meristem development continues in the notch, producing a complex, mature meristem with only left-right symmetry. This can be seen from changes in expression of genes like Mp*YUCCA2*, Mp*C3HDZ* and Mp*C4HDZ* within the apical notch and across the growing gemma (9, 24).

**Figure 1.**
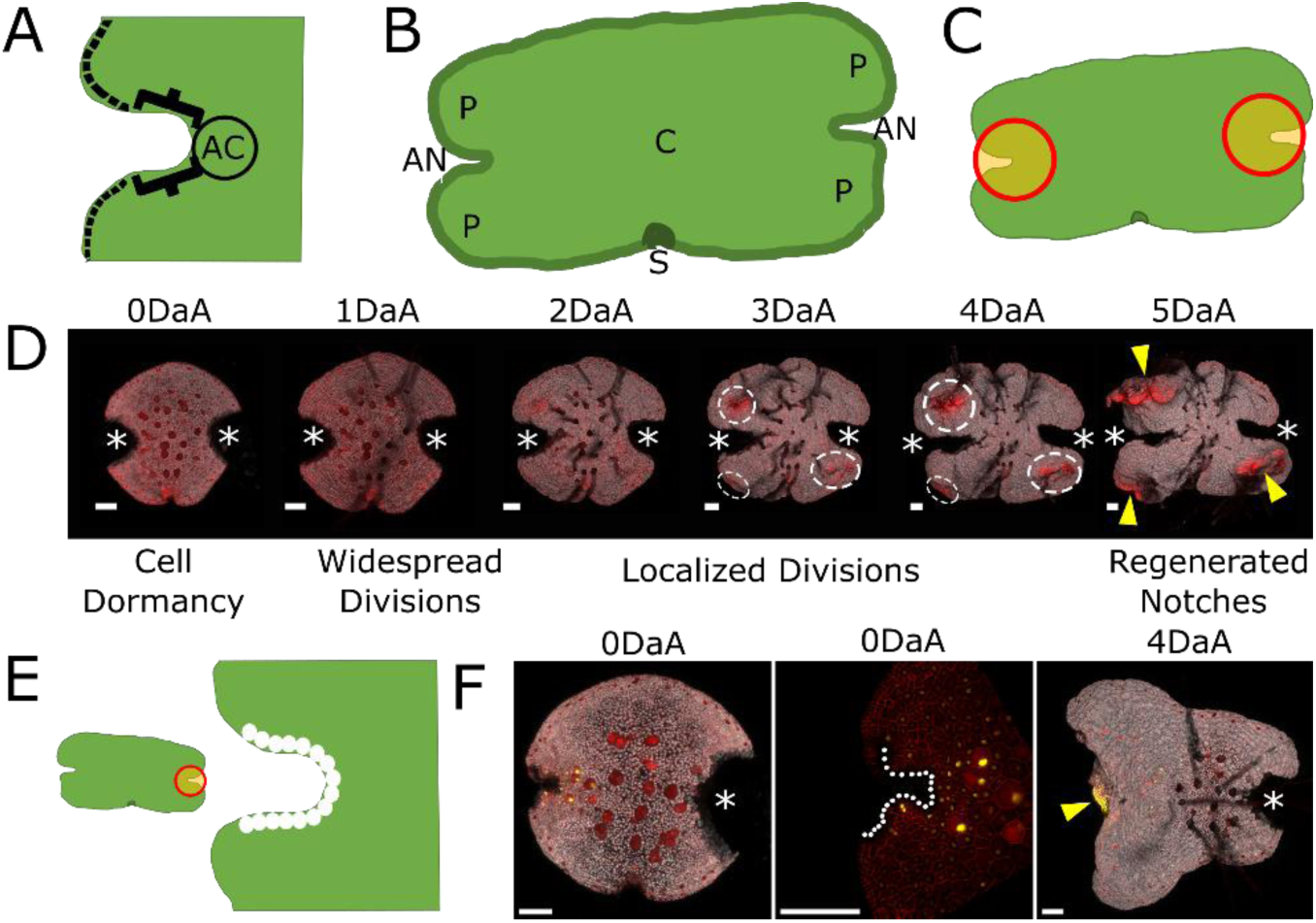
Laser ablation shows that the first row of cells in the apical notch are indispensable for meristematic activity. (A) Apical notch schematic. AC marks the apical cell at the notch apex (centre), flanked by subapical cells (brackets). These cells comprise the first row of cells. Further outwards the first row of cells is margin tissue (dashed line) (9). (B) Gemma schematic showing two apical notches (AN) at either end, the peripheral regions (P) to the sides and the central region (C) behind the notches. The stalk scar (S) is where the gemma was attached to the parent thallus within the gemma cup. (C) Schematic of the ablation pattern (orange bounded by red) for complete excision of all notches. (D) Time course of a gemma fragment expressing an mScarlet cell membrane marker (p5-UcE2:mSc-Lti6b) following complete notch excision. The four main phases of meristem regeneration are described. Excised notches are marked by asterisks. Cell division can be seen by increased density of red signal, localized patches of cell division are marked by dashed circles and regenerated notches marked with yellow arrows. Images were acquired at the indicated time-points after apical notch removal (days after ablation, DaA). (E) Schematic of the ablation patterns used in the fine scale gemma ablation experiment. Orange bounded by red denotes excised notch, white circles indicate ablated cells. (F) shows, from left to right, the 0DaA gemma (asterisk marking excised notch), 0DaA notch close up (ablated cells marked by white circles) and the 4DaA gemma (yellow arrow marks regenerated notch). Ablating the first row of cells in the apical notch is sufficient to induce meristem regeneration, which occurs just behind the ablated notch. Gemma shown is from the enhancer trap apical notch/meristem marker line ET239-P153. See SI Appendix Fig. S2*B* for the full time series. Confocal images show membrane marker in red, nuclear mVenus enhancer trap signal in yellow and chlorophyll autofluorescence in grey. Scale bars= 100μm.

The gametophyte apical notch meristem demonstrates apical dominance dynamics and lateral inhibition (25). Apical notches prevent the emergence of new active meristems (outside of dichotomous branching) and repress dormant meristems, such as gemmae within the gemma cup (26) or shaded apical notches (21). If active apical notches are removed, then meristems kept dormant by lateral inhibition start growing (27). If all apical notches are removed then apical dominance is lifted and meristems regenerate *de novo* (8, 11). Auxin has long been known to be involved in meristem regeneration in Marchantia (25, 27–30). Auxin is produced in the apical notch and is exported to the rest of the thallus, where it inhibits formation of new meristematic regions (i.e., apical dominance). Removing apical notches removes these auxin sources. Initially auxin levels drop in the remaining plant, and then rise as new meristems emerge (8).

Developing better insights into how meristems function is key to understanding plant development and producing beneficial new morphologies (2, 3). Understanding the molecular mechanisms of cellular reprogramming to convert different cell types into pre-meristematic and meristematic tissue is important in vegetative propagation of agricultural crops and offers an avenue to develop new transgenic methods (7, 31). Considerable progress on understanding apical meristems has been made from study of angiosperms, particularly *Arabidopsis* (32), however these systems have limitations, such as difficult access for imaging (particularly live imaging) (33) and the complexity of the gene networks involved (2, 4).

Although the pre-meristem and meristem of the Marchantia apical notch has received relatively little attention compared to angiosperm meristems (34, 35) it offers a promising opportunity to gain insights into meristematic tissues generally. The key advantage of Marchantia as an experimental system for plant biology is its streamlined nature. For many gene families the Marchantia genome has fewer copies compared to other model plants like Arabidopsis or Physcomitrella (34, 36), notably the minimal set of components related to auxin biosynthesis, transport and signalling (36–41). It is highly experimentally tractable, with fast growth rates and well-established transgenic protocols (42, 43). The gemma is well-suited for developmental experiments, since it grows in a stereotypical pattern. Crucially, gemmae have open development where almost all of the cells and tissues are easily accessible for microscopy without the need for dissection, sectioning or clearing (42).

Marchantia, as a bryophyte, occupies an important position in the land plant phylogeny (15, 44, 45). Although the thalloid gametophyte is not homologous to the dominant diploid sporophyte of vascular plants, phylogenetic analyses and molecular research finds that the apical notch (pre-)meristem shares many regulatory modules in common with angiosperm apical meristems (14, 46–51). The SAM model is thought to be how meristems operated in the first land plants (52), making findings from Marchantia applicable to all plant meristems (53). Inter-and intra-apical notch signalling in the Marchantia gemma allows study of pre-meristem and meristem communication, and of the gene networks regulating meristem morphology (8, 46, 47, 54, 55).

Meristem regeneration in gemmae provides an opportunity to investigate cellular reprogramming and how emerging pre-meristematic regions interact. This can provide insights into how Marchantia specifies new meristems during spore germination (9, 20, 56), gemma cup development (22) and apical notch bifurcation (21, 46), processes relevant to vegetative reproduction and lateral branching in crop species. In addition to its importance in plant evo-devo studies, Marchantia holds great promise as a chassis for synthetic biology (42, 57), and greater understanding of Marchantia meristems will further this research.

I used laser ablation and transgenic marker lines to make precise disruptions in the apical notches of gemmae and observed how this altered cell proliferation and apical dominance (56, 58, 59). My results point towards a model with a dynamic population of stem cells communicating within the apical notch and with the rest of the gemma. Only certain regions of the gemma can conduct these communication signals, and a balance of auxin sources and sinks is required to sustain meristematic activity. Disruption of this communication and balance induces new meristems to emerge, or causes existing apical notches to cease cell division.

## Results

### Laser ablation of the apical notch induces meristem regeneration

Marchantia plants expressing a plasma membrane marker fused to mScarlet driven by a ubiquitous promoter (p5-UcE2:mSc-Lti6b) allowed the visualization of new cell membranes and cell division. Laser ablation microscopy was used to remove both notches of a 0 days post germination (dpg) gemma (Fig. 1*C*) and the membrane marker line used to track meristem regeneration (Fig. 1*D*). This background experiment confirmed previous findings (60), that there are four distinct phases of regeneration in Marchantia. In the first ∼24 hours little or no cell division occurs. Between 24-48 hours after ablation there is rapid, widespread cell division across the gemma. After ∼48 hours cell division becomes restricted to localized patches of cell division (dashed circles). Approximately 5 days after ablation (DaA) meristem regeneration has progressed to formation of apical notches (yellow arrows).

A collection of enhancer trap lines were used to monitor apical notch activity and regeneration. The genes marked are not yet identified but these lines were all previously established to mark the apical notch/meristem region via nuclear mVenus signal (9) Control ablation experiments demonstrated that the location and progress of meristem regeneration could be monitored using the enhancer trap signal (SI Appendix Fig. S1). Nuclear mVenus marks patches of localized cell division, only persists in actively dividing regions and is maintained in active apical notches. Loss of marker signal indicates loss of apical notch identity and cell proliferation activity.

Examination of multiple gemmae with both notches completely removed (ablation pattern in Fig. 1*C*) did not find any clear predictability as to which regions would go on to form new apical notches, nor how many patches of localized cell division or regenerated notches there would be. The exception to this is the stalk scar region, which was never observed to initiate cell divisions.

Laser ablation microscopy can make fine scale perturbations in the apical notch to see how much tissue must be removed to induce meristem regeneration. One apical notch was entirely excised and in the other notch a number of rows of cells were ablated. Ablating one cell row deep into the apical notch was sufficient to induce meristem regeneration (Fig. 1*E*, *F* and SI Appendix Fig. S2*A*, *B*). Notch organisation was disrupted following ablation, with the area expanding outwards and meristem regeneration always occurring in the region behind the ablated row of cells (yellow arrow, Fig. 1*F*, SI Appendix Fig. S2*B*). Ablating the first three rows of cells was required for meristem regeneration to be induced elsewhere in the gemma, away from the notch (SI Appendix Fig. S2*C*, *D*). These background investigations established the first row of apical notch cells as the focus for subsequent laser ablation experiments.

### Laser ablation demonstrates interactions between apical notches

The current Marchantia literature centres around a model where, following germination, each apical notch contains a single apical cell. This divides, giving rise to lateral subapical cells, which comprise the rest of the first cell row in the apical notch. However until recently there has been little experimental detail on how these cell types are specified, how they interact with each other or with other apical notches (55, 56). I interrogated this in more detail by ablating specific parts of the crucial first row of apical notch cells in the developing pre-meristem. A flow chart of the outcomes of these experiments is provided in SI Appendix Fig. S3.

Ablating one side of the apical notch did not disrupt meristematic activity (i.e., cell proliferation and apical dominance), irrespective of whether or not there was another intact apical notch present in that gemma (SI Appendix Fig. S4*A*, *B*). However, leaving only a few apical notch cells intact on one side disrupted meristematic activity. If the other apical notch in the gemma was intact (SI Appendix Fig. S4*C*) then the partially ablated notch became inactive due to apical dominance from the intact notch. If the other notch was excised (SI Appendix Fig. S4*D*), the remaining cells in the partially ablated notch were insufficient to sustain apical dominance, and meristem regeneration occurred elsewhere in the gemma fragment. When a few apical notch cells were left intact on both sides of the notch then this partially ablated notch always lost meristematic activity (SI Appendix Fig. S4*E*, *F*). Notches with multiple small subpopulations of intact cells lost meristematic activity, even if there were more intact cells in total compared to functional notches where the intact cells comprised a contiguous row.

When both sides of the apical notch were ablated, but a few cells at the apex (including the presumed apical cell) were left intact, then the outcome depended on the status of the other notch in the gemma. If the other notch was excised (Fig. 2*A*), then the intact apex continued to function as an apical notch (blue arrow) and retained apical dominance as no new patches of cell division emerged elsewhere. When the other notch in the gemma was intact (Fig. 2*B*), then the intact apex alone could not sustain meristematic activity (grey arrow), even if more cells remained compared to cases where meristematic activity continued with the other notch excised (e.g., Fig. 2*C*).

**Figure 2.**
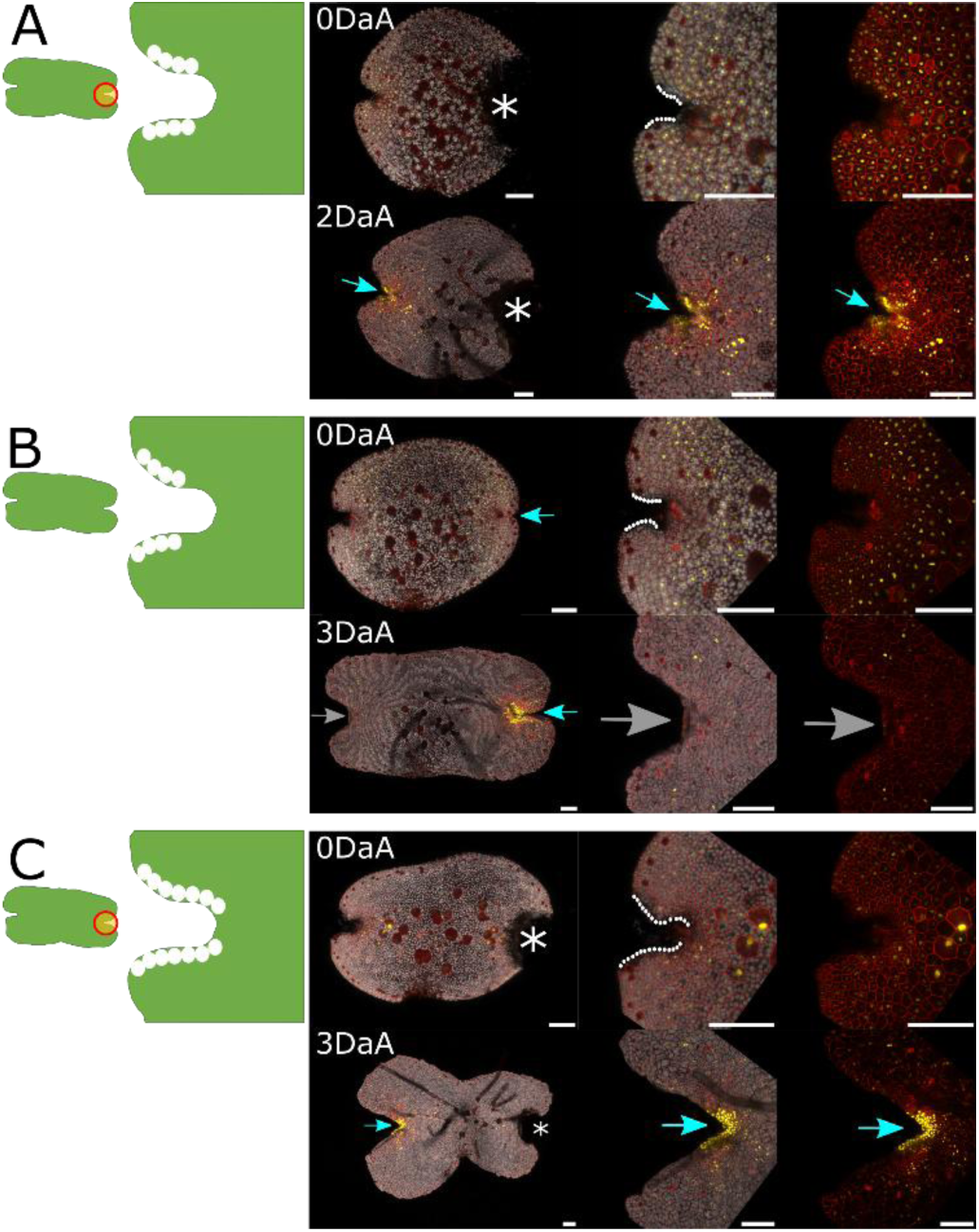
The notch apex, containing the apical cell, can sustain meristematic activity only if the other apical notch is excised. Ablation of the notch sides, leaving the notch apex (including the apical cell) intact, did not disrupt meristematic activity if the other apical notch in the gemma was excised (A), but when the other apical notch was intact then meristematic activity ceased (B). (C) Meristematic activity continued even if few cells were left intact at the apex if the other notch was excised. In each sub-figure a schematic of the ablation pattern is shown with white circles marking the location of ablated cells and orange bounded by red denoting excised tissue. Time courses are in the format whole gemma, notch close up, notch close-up without chlorophyll autofluorescence channel. Asterisks mark excised notches, blue arrows denote active apical notches, grey arrows mark apical notches that have lost meristematic activity. In the 0DaA images, white circles mark ablated cells. The enhancer trap apical notch/meristem marker lines used were ET239-P21 (A, B) and ET239-P153 (C). Scale bars=100µm.

Parts of the notch spanning the apex and sides were ablated, including where the apical cell is located (Fig. 3). When some of the apical notch cells were ablated, but a sufficient subpopulation remained as a contiguous row, then not only did the notch continue proliferative activity (blue arrows), but over the next 48 hours its growth pattern changed. By 3DaA, what was previously the centre of the intact subpopulation of apical notch cells has become the notch apex, i.e., the notch had reorientated. Ablation experiments on Mp*YUC2* promoter marker line gemmae (Fig. 3*C*) found that promoter activity continued during reorientation, and that at 3DaA the reorientated apex was the focus of Mp*YUC2* expression, and therefore auxin biosynthesis. This shows that reorientation is not just superficial, but that the mechanistic activity of the apical notch also shifts position.

**Figure 3.**
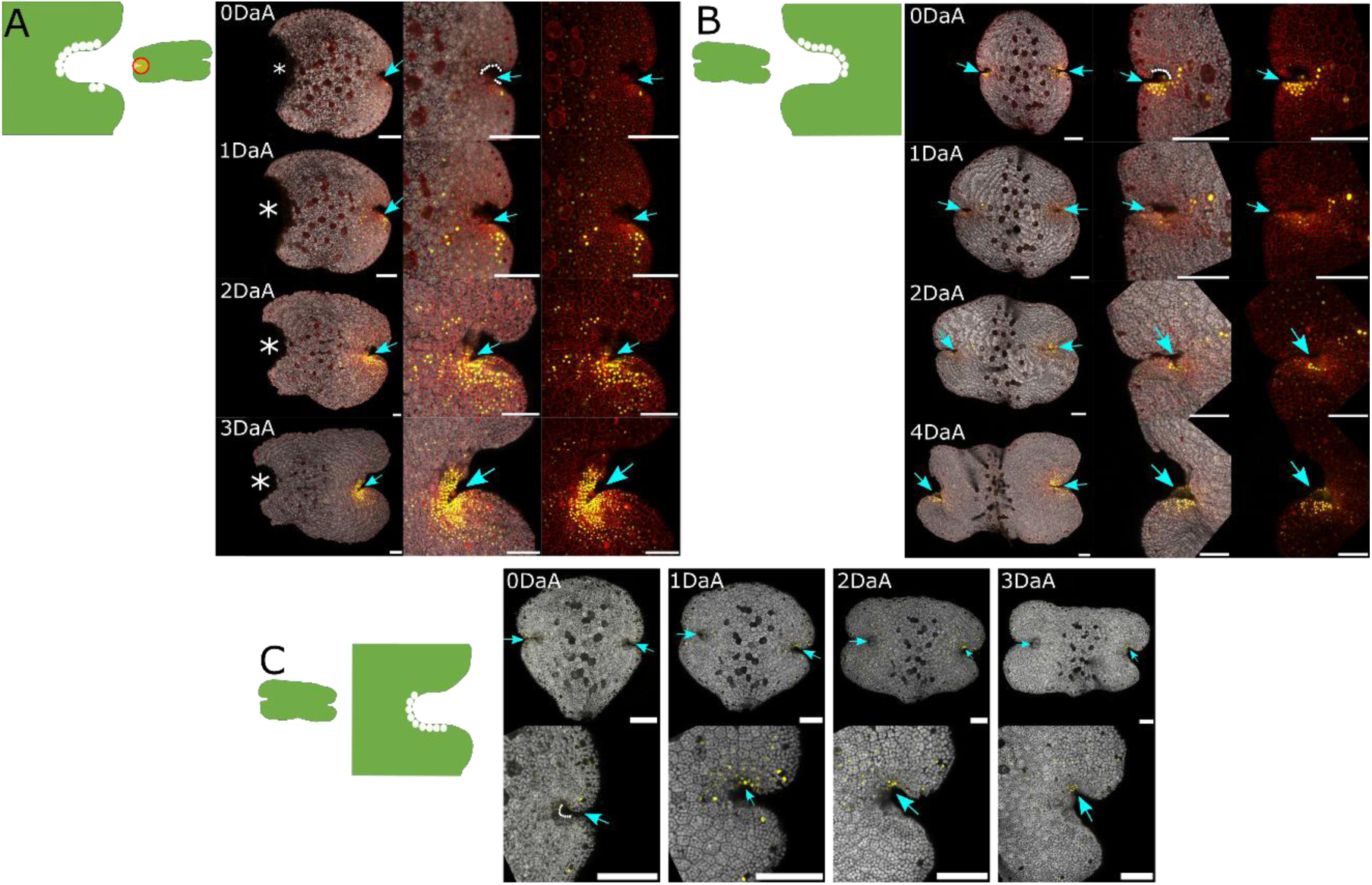
Notch reorientation can occur after partial ablation of the notch apex, independent of the status of the other apical notch in the gemma. Partial notch ablation, such that the intact stem cell subpopulation spans the apex and one side of the notch, results in reorientation of the apical notch. The notch changes shape, forming a new apex whose location corresponds to where the centre of the subpopulation of intact cells was originally. Reorientation occurs whether the other notch in the gemma was excised (A) or intact (B, C). Meristematic activity continued during reorientation, as indicated by mVenus expression in enhancer trap lines (A, B). Mp*YUCCA2* promoter activity (C) shows that the new apex becomes the focus of auxin production in the reorientated notch. In each sub-figure a schematic of the ablation pattern is shown with white circles marking the location of ablated cells and orange bounded by red denoting excised tissue. Time courses in A and B are given in the format whole gemma, notch close up, notch close-up without chlorophyll autofluorescence channel. Asterisks mark excised notches; blue arrows denote active apical notches. In the 0DaA images, white circles mark ablated cells. A and B show the enhancer trap apical notch/meristem marker lines ET239-P21; C shows a Mp*YUC2* promoter marker line with mTurquoise signal in yellow. Scale bars=100µm.

Reorientation occurred irrespective of where the apex of the original notch had been before ablation and did not appear to be influenced by whether the original centre of the notch (i.e., the location of the apical cell) was ablated or not. During the first 48 hours after ablation, no new regions of cell division appeared (if the other apical notch in the gemma had been removed), nor were reorientating notches subject to apical dominance-related inhibition (if the other notch in the gemma was intact). There were no changes in Mp*YUC2* promoter activity in the partially ablated notch or across the rest of the gemma that might indicate an interruption or major spatial reorganization of auxin biosynthesis. Notch reorientation occurred even when gemmae were grown on media containing 3µM 1-naphthylacetic acid (1-NAA) (SI Appendix Fig. S5), a pharmacological inhibitor of regeneration (8). This indicates that reorientating notches continued to be apically dominant, and that the morphological changes involved in notch reorientation were not due to *de novo* meristem regeneration.

### Meristem regeneration involves communication between regions of cell division

Fig. 4 and SI Appendix Fig. S6 show a gemma ablated so that one apical notch was entirely excised and in the other notch a short, contiguous subpopulation of cells was left intact. Initially, the rest of the gemma was released from apical dominance and proceeded to the early phases of regeneration. By 2DaA there were regenerating patches of localized cell division with dense enhancer trap marker signal (dashed circles, Fig 4*C*). Simultaneously, the partially ablated apical notch also recommenced cell division (white arrow, Fig. 4*C*). By 3DaA this region was the most prominent area of marker signal, though other patches of signal were visible (SI Appendix Fig. S6*D*). Five days after ablation (Fig. 4*D*, SI Appendix Fig. S6*F*), the notch that was originally partially ablated had the characteristic apical notch morphology and was actively proliferating (blue arrow). It had re-established apical dominance, and the other patches stopped division and meristem regeneration (indicated by loss of marker signal, green arrows). However, these areas progressed considerably through meristem regeneration, with sufficient cell divisions to produce protrusions on the thallus.

**Figure 4.**
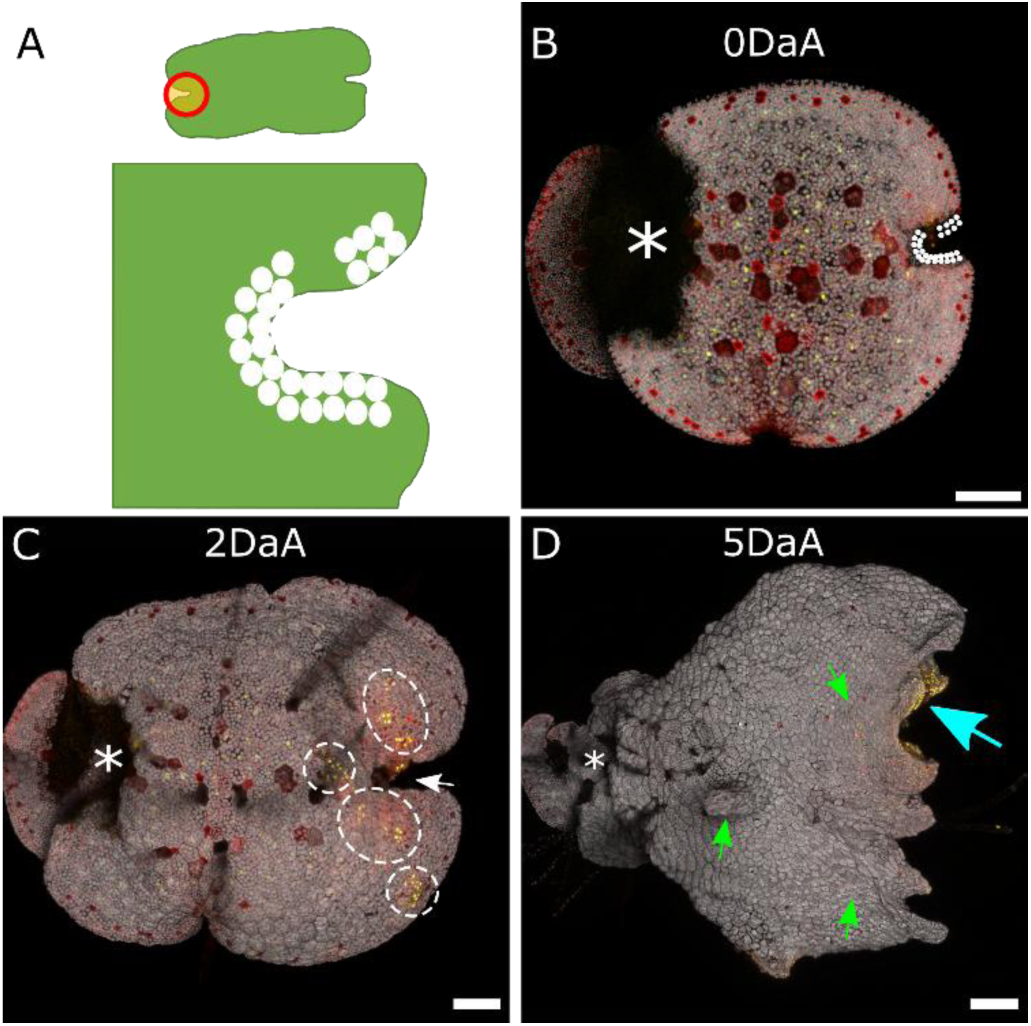
Time course demonstrating re-establishment of apical dominance by a partially ablated apical notch. (A) Schematic of the ablation pattern used. White circles mark ablated cells, orange bounded by red marks excised tissue. (B) shows the gemma immediately after laser ablation. White circles mark the position of ablated cells, asterisk indicates the entirely excised apical notch. (C) At 2DaA patches of localized cell division and marker signal appeared (dashed circles), even though the original, partially ablated notch still displayed marker signal (white arrow). (D) By 5DaA, the original partially ablated notch had meristematic activity (blue arrow), as indicated by the presence of dense marker signal, and the apex of this notch had reorientated. The areas that were patches of cell division had no marker signal (i.e., no cell division activity) and remained as protrusions on the gemma surface (green arrows). The complete time course is given in SI Appendix Fig. S6. The enhancer trap apical notch/meristem marker line used was ET239-P125. Scale bars=100µm.

### Directional flow of signal out from the apical notch causes apical dominance

The apical dominance signal produced at the apical notch must flow outwards in order to influence the rest of the gemma. Tissue was ablated in different orientations to the apical notch to see if the resulting incisions could interrupt the flow of signal. Incisions made perpendicular to the notch had no obvious effect (SI Appendix Fig. S7*A-E*). Incisions made parallel to the notch (SI Appendix Fig. S7*F-O*) resulted in more cell division proximal to the incision and increased cell expansion distal to the incision. This effect was not seen when the portion of the apical notch immediately proximal to the parallel incision was ablated; instead cells on both sides of the incision ceased dividing and expanded.

More extensive incisions found that the type of connection to the rest of the gemma was crucial for transport of the apical dominance signal (Fig. 5). Connections formed of tissue from the central part of the gemma (Fig. 5*A-C*) communicated the repressive signal from the active apical notch (Fig. 5, blue arrows) and did not see meristem regeneration elsewhere. This was the case even when the connecting tissue was only a few cells wide.

**Figure 5.**
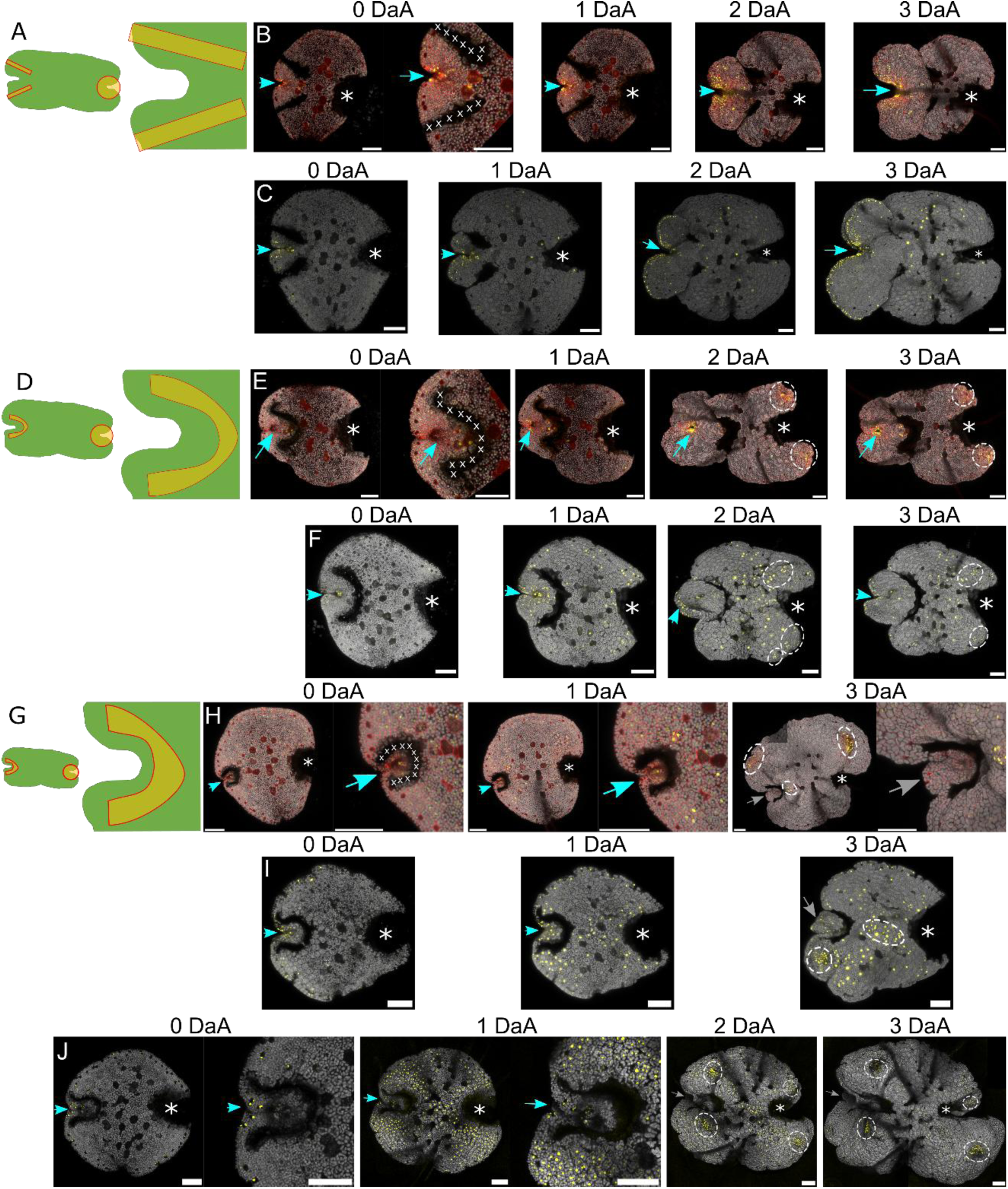
Ablating tissue around the apical notch demonstrates that central regions of the gemma can transport apical dominance signal, but peripheral regions cannot conduct apical dominance signal. (A) Schematic of the ablation pattern used in time courses (B) and (C). The intact apical notch maintained meristematic activity (blue arrows). No meristem regeneration occurred in the gemma fragment attached to this notch by a central connection. (C) shows a marker line for Mp*PIN1* promoter activity. There is signal at the notch and the gemma edge but little in the centre of the gemma, and none in the central connection itself. (D) Schematic of the ablation pattern used in time courses (E) and (F). The apical notch remained active (blue arrows) however meristem regeneration (dashed circles) occurred in the gemma fragment attached to this notch by the peripheral connections. In (E) this was marked by the appearance of new enhancer trap mVenus marker signal, while in (F) this was indicated by proliferation of Mp*PIN1* promoter signal throughout the gemma, and later by clusters of Mp*PIN1* promoter signal. Though signal was visible in the peripheral connections early on in time course (F), this connection apparently did not conduct apical dominance signal from the intact notch. (G) Schematic of the ablation pattern used in time courses (H)-(J), where the gemma was ablated to leave only a small amount of tissue surrounding the intact apical notch. By 3DaA, regenerating meristems emerged (dashed circles). The intact apical notch was initially active, indicated by enhancer trap marker expression (blue arrows in (H)) but by 3DaA this notch was inactive (grey arrows), with the cells losing marker signal, ceasing division and expanding. In the Mp*PIN1*(I) and Mp*YUC2*(J) promoter marker line time courses, loss of signal was observed at the inactive notch (grey arrows). At 1DaA there is widespread Mp*YUC2* promoter activity across the gemma fragment, but this becomes restricted to progressively smaller regions as meristem regeneration proceeds (J). Asterisk marks entirely excised notch, blue arrows denote active apical notches, grey arrows denote inactive apical notches, dashed circles mark patches of localized cell division. In schematics (A, D, G) X marks incision ablations, orange bounded in red denotes excised tissue. Gemmae shown are from the enhancer trap apical notch/meristem marker line ET239-P153 (B, E), ET-239-P125 (H), Mp*PIN1* promoter marker line with mVenus signal in yellow (C, F, I) or Mp*YUC2* promoter marker line (J). Scale bars= 100µm.

In contrast, connections formed from tissue around the gemma periphery (Fig. 5*D-F*) did not communicate the repressive apical dominance signal. Multiple regenerating meristematic regions emerged in the rest of the gemma, despite there being a fully functioning apical notch present. If the region around the original apical notch included significant amounts of surrounding tissue (Fig. 5*E,F*), then the apical notch continued to grow and develop as normal (blue arrows), even as meristem regeneration occurred elsewhere (dashed circles).

If this region contained just the apical notch and cells immediately surrounding the notch, then the outcome was different (Fig. 5*G, H*). The rest of the gemma was released from apical dominance and meristem regeneration proceeded (dashed circles). The original notch continued meristematic activity for the first day after ablation, indicated by enhancer trap marker signal persisting. At 2DaA this signal faded and cell division ceased, and by 3DaA the notch was inactive (grey arrows). This occurred in cases where the apical notch and immediately surrounding region was totally disconnected from the rest of the gemma (SI Appendix Fig. S8).

This observed self-inhibitory feedback is characteristic of auxin (37, 55, 61). A reporter line marking activity of the canonical polar auxin transporter Mp*PIN1* promoter (38, 45) shows that in 0dpg gemmae signal is focused around the apical notches, is present around the gemma edge and in occasional cells across the gemma centre. This supports findings that the apical notch is an auxin source (55), and suggests that auxin transport and signalling is more widespread. Marker line gemmae were ablated according to the incision patterns described above (see Fig. 5*C, F, I*). Fig. 5*C* showed no expression central tissue connection, yet the intact notch retained apical dominance over the rest of the gemma. Conversely, Fig. 5*F*, *I* show that peripheral connections had Mp*PIN1* promoter activity, yet meristem regeneration occurred elsewhere in the gemma (dashed circles). The reduced Mp*PIN1* promoter signal around the apical notch by 3DaA in Fig. 5*I* reflected the loss of meristematic activity in that notch (grey arrow). This also occurred in Mp*YUC2* marker line gemmae (Fig. 5*J*), where there was loss of Mp*YUC2* promoter activity in the isolated notch as it ceased proliferation and the cells expanded. Elsewhere there was widespread Mp*YUC2* promoter activity at 1DaA, indicating reorganization of auxin biosynthesis activity due to falling auxin levels in the rest of the gemma. Mp*YUC2* promoter activity then became progressively more focused to smaller areas as meristem regeneration proceeded.

### Larger gemma fragments regenerate meristems faster than smaller gemma fragments

Since the size of the surrounding region has an effect on apical notch activity, I examined how the size of the gemma fragment influences meristem regeneration. A standard-sized area around the apical notch region was entirely excised in a number of gemmae from the enhancer trap lines ET239-P125 and ET239-P33. The remaining gemma fragments were then ablated to leave a circle of tissue with a diameter 40, 60, 80 or 100% the notch-to-notch distance of the original gemma. The reappearance of the enhancer trap marker signal and the appearance of notch morphology was used to measure the speed of meristem regeneration (Fig. 6, see SI Appendix Fig. S9 for regeneration timings and SI Appendix Fig. S10 for a representative set of images from the time course). The timings of when patches of localized cell division with marker signal appeared (dashed circles, SI Appendix Fig. S10), and when notch-type morphology re-formed (yellow arrows, SI Appendix Fig. S10) were recorded (Fig. 6 and SI Appendix Fig. S9).

**Figure 6.**
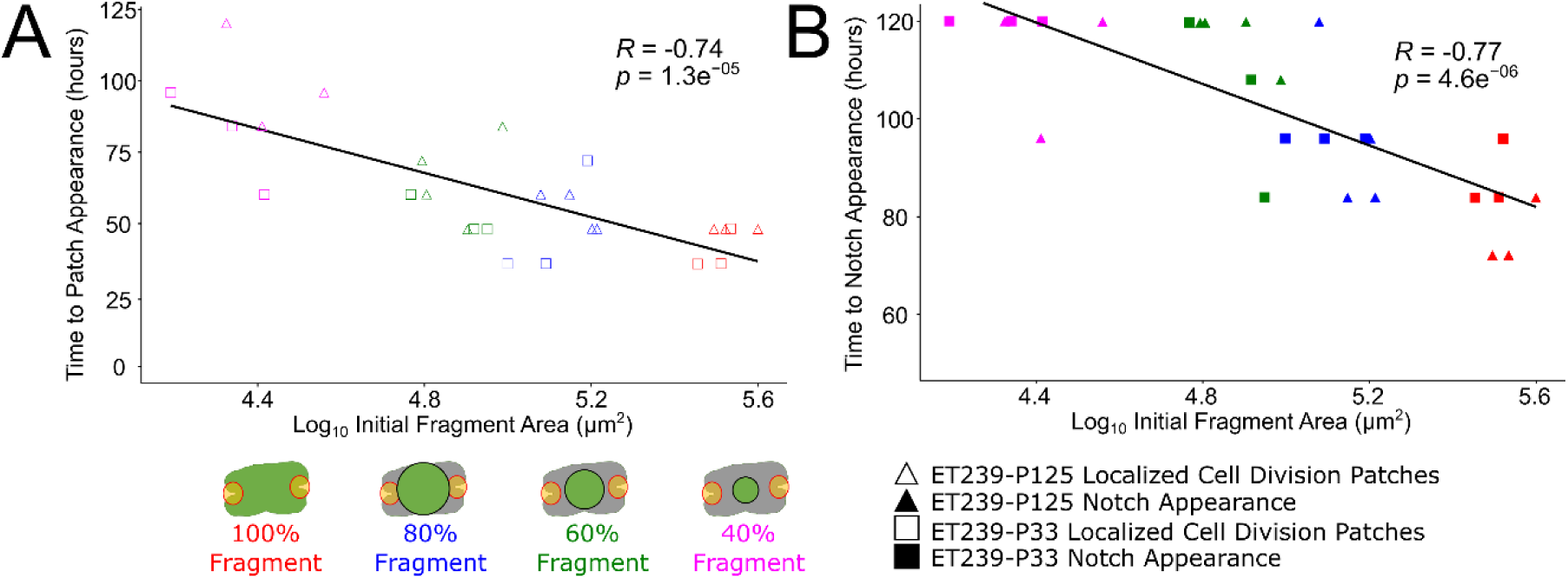
A 108-hour time course using two different enhancer trap apical notch/meristem marker lines shows that larger gemma fragments have faster emergence of patches of localized cell division and faster appearance of morphologically recognizable notches. Plotting log_10_ fragment area versus time to appearance of localized patches of cell division (A) or notch morphology (B) finds a significant Pearson’s correlation coefficient between fragment size and regeneration speed. Where patches or notches had not appeared within the duration of the experiment, time was given as 120 hours. A schematic of the ablation pattern used is given at the bottom, with orange bounded by red indicating an entirely excised apical notch. The black circle shows the laser ablation circle trace used, with the circle diameter calculated as a percentage of the original notch-notch distance. Open symbols denote appearance of patches of localized cell division with enhancer trap marker signal; closed symbols denote when notch morphology within a dense patch of marker signal was first observed; symbol colours relate to the percentage fragment size as indicated in the schematics. See SI Appendix Fig. S9 for regeneration timings and S10 for a representative set of time course images.

Over the 108-hour time course it was observed that larger fragments had patches of marker signal reappear significantly earlier (Fig. 6*A*) and had significantly faster regeneration of apical notch morphology (Fig. 6*B*) compared to smaller fragments.

## Discussion

### Laser ablation of transgenic Marchantia lines reveals minimum requirements for meristem regeneration

Previous experiments investigating Marchantia regeneration were limited to surgical manipulations on older thalli, removing all of the apical notch with scalpel blades (8, 11, 12, 27). These cutting experiments demonstrated that only when all apical notches were removed from a thallus would new meristematic regions emerge. Laser ablation microscopy allows for finer scale disruptions of the apical notch pre-meristem of 0dpg gemmae, which is more accessible for ablation than the fully mature meristems of older gemmae. Laser ablation demonstrated that removal of the first row of cells in the apical notch is sufficient to induce meristem regeneration. It reveals a previously unappreciated substructure within the pre-meristem of the 0dpg gemma (Fig. 7*A*), with the immediate first row of apical notch stem cells being critical for (pre-)meristem maintenance and the second and third rows being “primed” for meristem regeneration in preference to the rest of the gemma. The cells in these rows are divisionally active and have the Mp*ARF2*-related gene expression networks of early ontogeny (55), therefore less reprogramming would be involved to acquire meristem identity compared to other cells.

**Figure 7.**
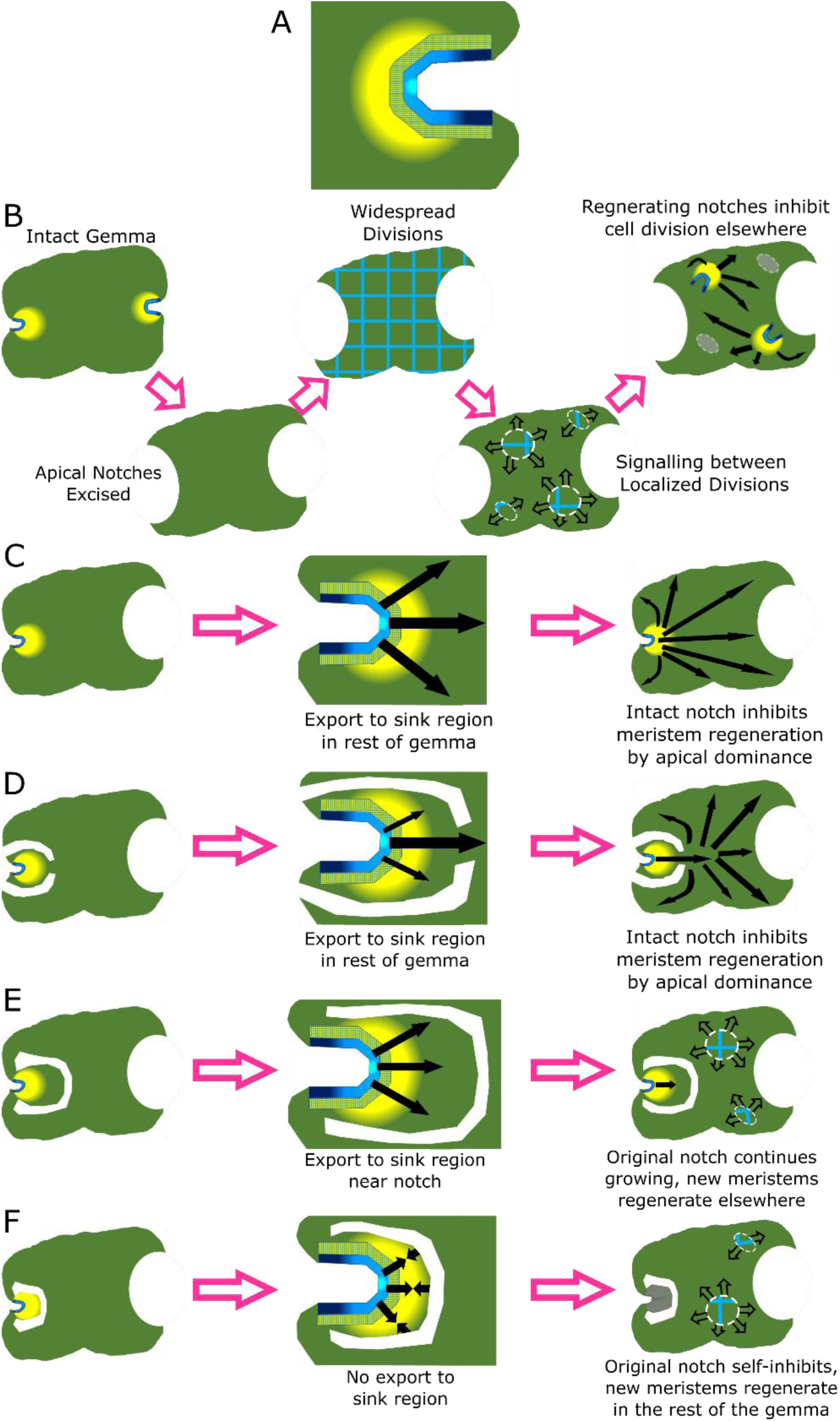
A model of apical notch organization, meristem regeneration and apical dominance signalling in Marchantia. (A) The stem cells in the first row of the apical notch (in blue) are critical for (pre-)meristem maintenance. The cells at the centre of this population are distinct from the cells towards the edge (blue gradient). The cells in the two rows behind are primed for meristem regeneration (blue hatching). The stem cells emit a signal regulating cell division and expansion in the surrounding region (yellow gradient). (B) Process of meristem regeneration after excision of both apical notches. Initially there are widespread cell divisions (blue grid), then cell divisions are restricted to localized patches (circles) which communicate (open arrows). Eventually one or more of these patches passes a developmental threshold and produces a repressive apical dominance auxin signal (black arrows) that inhibits cell division elsewhere. (C) When one apical notch is intact, it emits an apical dominance auxin signal (black arrows) that prevents the emergence of any new meristems. (D) When one apical notch is intact and linked to the rest of the gemma fragment through a central connection, then apical dominance signal is exported (black arrows) and no new meristems regenerate. (E)-(F) When one apical notch is intact and linked to the rest of the gemma fragment through a peripheral connection, no apical dominance signal is exported (black arrows), and meristem regeneration occurs elsewhere in the gemma fragment (circles). If the intact apical notch is surrounded by sufficient auxin sink tissue then it continues meristematic activity (E). If the intact apical notch is not surrounded by sufficient sink tissue, then auxin builds up and this notch becomes inactive (marked in grey) due to self-inhibition (F).

Beyond this there appears to be no wider predictability as to which regions will regenerate new meristems (Fig. 7*B*). The exception is the stalk scar region, which was never observed to regenerate new meristem. These are the oldest cells in the gemma, produced at the initiation of gemma formation within the gemma cup (18, 22). It may be the case that, just as the youngest cells in the gemma are primed for meristem regeneration, the oldest cells are excessively complex to reprogram. This regenerative flexibility was reported in early studies (12), and is in contrast to vascular plants, where in the post-embryonic plant only certain somatic cell types can form new meristem. It also differs from cutting experiments done on older thalli, where regeneration always occurred at the apical midrib on the basal explant (8, 11).

### Apical notch organization and communication

The standard model of the Marchantia meristem involves a single pluripotent apical cell with four division planes, located at the centre (apex) of the notch. This apical cell divides to form dorsal, ventral and lateral merophytes/sub-apical cells, which then go on to form the various tissues of the thallus (18, 19, 34). The apical cell in the 0dpg gemma pre-meristem has primarily been defined by morphological characterization rather than direct experimental evidence (18, 24, 47). For dichotomous branching and apical notch bifurcation, two apical cells must be present (46, 47). There is debate over the bifurcation mechanism: *de novo* formation of a new apical cell at a distance from the original one; loss-of-fate of the original apical cell and *de novo* formation of two new apical cells some distance from each other; or division of the original apical cell to produce two new apical cells, which each then divide, producing subapical cells between them and displacing the two apical notches apart. In more complex plant meristems (e.g., angiosperm SAM) there is a population of pluripotent cells that divide and maintain the meristematic stem cell niche (32).

I tested how this model applies to the pre-meristem of the 0dpg apical notch experimentally, by ablating portions the crucial first cell row of the apical notch (i.e., the apical cell and subapical cells). This showed that the important factor is the number of intact cells in this row that are in contiguous contact. The notch apex (containing the apical cell) seemingly has a greater functional capacity than the cells on the sides of the notch, since this central subpopulation alone could sustain meristematic activity if no other apical notches were present in the gemma. This confirms communication between apical notches within a gemma.

When the first row of apical notch cells was ablated to remove one side and part of the notch centre, there was a sufficiently large contiguous subpopulation of cells to sustain cell proliferation and apical dominance (Fig. 3). After 4 days the apical notch had reorientated itself, with a new apex where the centre of the intact contiguous subpopulation of cells was originally. Apical notch mechanistic processes also shifted, shown by the focusing of Mp*YUC2* promoter activity around the reorientated notch apex (Fig. 3*C*). This was not *de novo* meristem regeneration of a new apical notch; instead it appears to be a case of remodelling, where divisions of the subapical cells create a new apex, presumably with an apical cell at its centre. This happens even under conditions where meristem regeneration cannot occur, for example if another apical notch is present in the gemma, or in the presence of 1-NAA, an inhibitor of meristem regeneration. That apical notch reorientation (Fig. 3) is not *de novo* meristem regeneration is supported by the fact that no new patches of cell division emerge elsewhere in the gemma (*c.f.* Fig. 4). It also suggests that the reorientation process is to some extent decoupled from auxin signalling, unlike *de novo* meristem regeneration.

These experiments demonstrate that there is a population of stem cells in the apical notch. What is important for cell proliferation and apical dominance is that there is a sufficient number or “quorum” of these cells intact and in contiguous contact. The characteristic notch morphology emerges from differential rates of cell division and expansion from the quorum centre to the edges (62). The new apex of the reorientated notch is at the centre of this quorum, not necessarily the cells that were closest to the original notch apex and apical cell. Under this updated model, the “Apical Cell” is not a permanent cell identity that is defined at gemma germination but instead emerges from the communication dynamics within the notch. It is this feedback process (46, 47, 50) across the entire stem cell population that sustains meristematic activity.

The requirement for contiguous contact means that cells within the apical notch must communicate directly. The importance of such paracrine signalling in Marchantia thalli has been noted previously, where disrupting plasmodesmata connections caused cells to independently regenerate new thalli (63). Paracrine signalling would provide a mechanism to regulate cell division rates from the apex to the sides of the notch, and offer a basis for dichotomous branching, as previously suggested by Hirakawa and colleagues (46, 47). The formation of two distinct, active meristematic centres could be closely regulated if they arise from one contiguous population of apical notch cells. These two apical notch meristems could co-exist in close proximity because each consists of an equally sized quorum. A quorum of dividing cells may also be required for the production of apical cells, for example in the sporeling protonema-to-prothallus transition (9, 20, 56).

### Partial apical notch ablation reveals that meristem regeneration is a flexible process

Multiple regions in a gemma can begin the regeneration process, but not all will eventually form apical notches (9), just as not all apical notches are proliferative (21, 26, 27). This is attributed to some regenerating regions having apical dominance or lateral inhibition effects (11). Ablation experiments (Fig. 4, SI Appendix Fig. S6) suggest that there are certain developmental stage thresholds that govern dominance during regeneration. The partially ablated notch reasserted apical dominance over larger patches of cell division, showing that developmental stage is more important than the size of the regenerating region.

Loss of meristematic activity can be reversed, and *de novo* meristem regeneration can be stopped even at a late stage. There may be a hierarchy of threshold stages during the processes of meristem formation and regeneration. This would explain the strong apical dominance effects of the apical notch pre-meristems in the germinating gemma, even before full meristem maturity. Since the partially ablated notch in Fig. 4 was already composed of stem cells it progressed faster to the key threshold stage for apical dominance. Regenerating regions that have passed specific thresholds could dominate other, less mature regions, while multiple regions could co-exist if they develop simultaneously (8, 9, 46, 47). This could provide a checkpoint mechanism for meristem regeneration and prevent too many regions emerging in too close proximity, and explain how bifurcating meristems could co-exist in close proximity, if they co-ordinate development through threshold stages or are beyond a threshold stage at formation.

### Apical dominance signal transport across the gemma

The partial notch ablation experiment shows that apical dominance signalling in Marchantia is an active process that can be temporarily disrupted and re-established. Lasering incisions demonstrates that apical notches produce a signal that flows outward to influence the rest of the gemma (Fig. 7*C*). Incisions parallel to the notch (SI Appendix Fig. S7*F-O*) show more cell division proximal and more cell expansion distal to the incision. This was only observed when the portion of the notch closest to the incision was intact, irrespective of the integrity of the cells at the notch apex. Cells distal to the incision did not begin meristem regeneration, so while some signals flow directly from the notch, the apical dominance signal was transported around the parallel incisions.

Further experiments to test which regions conduct the apical dominance signal found a difference between connections formed from central and peripheral tissue (Figs. 5, 7*D, E*). This was not dependent on connection width, as central connections 1-2 cells wide did not see meristem regeneration, but wider peripheral connections did. This means that the apical dominance signal must be carried by a specific mechanism, and not all cells possess this mechanism. This may link to the observation that most auxin transport and meristem regeneration occurs along the ventral midrib of mature thalli (8, 11, 64), though gemmae have no midribs or defined dorsoventrality at germination (18, 24).

Auxin is the prime candidate for this apical dominance signal. Auxin regulates processes such as apical dominance (6) and meristem regeneration (7) in all plants, including Marchantia (28, 29). A reporter line marking promoter activity of the auxin biosynthesis gene Mp*YUC2* (26, 65) shows that at 0dpg there is signal at the apical notches and throughout the gemma, but this becomes more focused around the notch in older gemmae (9). This shows that auxin is produced in the apical notch and must be exported to the rest of the thallus, confirming recent findings by Flores-Sandoval and colleagues (55).

That study suggested that the canonical MpPIN1 is responsible for auxin transport immediately surrounding the apical notch but did not examine longer range auxin signalling across the gemma. Canonical PIN distribution (9, 38, 45) does not explain why an auxin-based apical dominance signal is exported through central connections but not peripheral connections. Mp*PIN1* promoter expression is high at the gemma edge, present in peripheral connections and absent from central connection cells (Fig. 5*C, F*). Furthermore, meristem regeneration still occurs in Mp*pin1* mutants, albeit more slowly (54). It may be the case that other auxin transporters (36, 38, 66) are involved. The Marchantia non-canonical short PINs are promising candidates as recent work found that they localize to the plasma membrane and are capable of auxin export.

MpPINW has an asymmetric distribution in the plasma membrane, and this asymmetry appears to point centrally rather than peripherally in cells around the apical notch (67). It is notable that meristem regeneration involves widespread Mp*PIN1* and Mp*YUC2* promoter activity, subsequently followed by marker signal concentrating around those regions of localized cell division. This means that meristem regeneration involves reorganization of the auxin transport machinery as well as auxin biosynthesis (dashed circles, Fig. 5*F, I, J*) (54).

### Balance between auxin sources and sinks is required for Marchantia meristem maintenance and regeneration

Auxin has a self-inhibitory effect on its own biosynthesis (37, 61) and elevated auxin levels prevent meristem regeneration (8, 25). In Marchantia, this self-inhibitory effect is believed to be mediated by auxin up-regulating Mp*ARF1* expression, which then antagonizes *MpARF2* expression and auxin synthesis (55). This means there must be balanced auxin transfer between sources, where auxin is produced, and sinks, where auxin is degraded (26, 65, 68). The MpR2DII degron reporter system has shown that the apical notch meristem is the auxin source while the surrounding tissue acts as an auxin sink (55).

If the auxin sink is too large compared to the source then apical dominance is lifted and new meristems emerge; if the sink is too small compared to the source then auxin build-up interferes with meristematic activity. In Marchantia, an example of the former is seen in the short-term response to partial apical notch ablation (Fig. 4). Examples of the latter are apical notches isolated by peripheral connections (Fig. 5*D-I*), or by complete separation from the rest of the gemma fragment (SI Appendix Fig. S8). Here, the cells immediately surrounding the apical notch are the only available sinks. When this region is large the isolated notch continues growing normally (Fig 7*E*), but when this is small the sink is insufficient, accumulating auxin self-inhibits and the apical notch ceases cell proliferation (Fig. 7*F*).

The auxin source-sink balance is also disturbed by apical notch removal. With no sources, auxin levels fall across the remaining fragment. Subsequently new sources (regions of cell division) emerge (Fig. 5*J*), until the balance is restored by the regeneration of apical notches (8). A hypothesis that emerges from this is that if there are a greater pool of cells to act as auxin sources or sinks, then regeneration will proceed faster compared to fragments with fewer cells. The results of the experiments presented in Fig. 6 (see also SI Appendix Fig. S9 and S10) support this. An extreme version of the relationship between fragment size and regeneration speed is the case of isolated thallus cells (9, 10, 69). Here it can take 5-7 days for cell divisions to occur, with recognizable notches only appearing after 14+ days. Marchantia meristem regeneration presents a simplified system to study auxin self-inhibition and the balance between auxin sources and sinks. This provides a basis for research into more complex scenarios, e.g., lateral organ formation (3, 6), wounding responses (58, 59).

## Conclusions

An outline model for the apical notch of a 0dpg Marchantia gemma is given in Fig. 7. A population of stem cells are found in the first row of the apical notch, behind which are 2 rows of cells primed to acquire stem cell identity. The centre of the stem cell population forms the apex of the notch by having lower rates of cell division compared to the edges. The stem cells produce intra-notch signals maintaining the quorum, short-range signals regulating cell division and expansion, and long-range signals sustaining apical dominance across the gemma. The apical dominance signal, likely auxin, is carried through the centre of the gemma and not the peripheral regions. If neither apical notch has a sufficient quorum of intact stem cells in contiguous contact, or if the apical dominance signal cannot flow to the rest of the gemma, then new meristems emerge elsewhere in the gemma.

## Materials and Methods

Vector constructs were built according to the Loop Assembly protocol (42, 70) from L0 parts, L1 and L2 plasmids in the OpenPlant Loop Assembly toolkit (42). Lines used for each figure are given in SI Appendix Supporting Text. Full plasmid sequences for vector constructs are given in SI Appendix Dataset S1. Plasmid vectors were transformed into Marchantia sporelings (Cam accession) by agrobacterium-mediated transformation according to previously described methods (42, 43). T0 sporelings and subsequent G1 plants were grown at 21°C under continuous light (intensity= 150 μmol/m^2^/s) on 1.2% w/v agar (Melford capsules A20021) plates of Gamborg B5 media with vitamins (Duchefa Biochemie G0210) prepared at half the manufacturer’s recommended concentration and adjusted to pH 5.8. The media was supplemented with 100µg/ml Cefotaxime (BIC0111; Apollo Scientific, Bredbury, UK) to reduce bacterial contamination, 20µg/ml Hygromycin (10687010; Invitrogen) for selection and 3µM 1-NAA (N1641, Sigma-Aldrich) for regeneration inhibition experiments.

Imaging was carried out using a Leica Sp8 confocal microscope or Zeiss LSM800 upright confocal microscope (SI Appendix Fig. S5). Laser ablation experiments were performed using a Leica LMD6000 laser dissection microscope system or Zeiss PALM MicroBeam Laser Capture Microdissection Microscope (SI Appendix Fig. S5). Microscopy settings are given in SI Appendix Supplementary Information Text and SI Appendix Supplementary Information Table 1. Images were processed using LasX, ZEN Blue and Fiji/ImageJ software packages. Figures were produced using the QuickFigures (71) and Stitching (72) Fiji plugins. Consistent growth and marker signal responses were observed in at least three gemma replicates taken from two different lines, except for Fig. 4/SI Appendix Fig. S6, where re-establishment of apical dominance by a partially ablated apical notch was observed once. Pearson’s correlation coefficient analyses were performed using R Statistical Software v4.5.2 (The R Foundation for Statistical Computing).

Propidium iodide (PI) staining was used to verify that laser ablation did not cause widespread stress or damage to the gemma. WT and enhancer trap line gemmae were incubated in 10µl of 100µg/ml PI (Sigma) on a glass coverslip for 10 minutes at room temperature, ensuring that the gemma was not mechanically damaged or desiccated during incubation. Staining, ablation and imaging was carried out in various orders to ensure that any PI signal relating to cellular damage was solely due to laser ablation. All experiments showed that the laser settings killed the targeted cells, cellular damage was localized to the ablated zone and there was no widespread cell death or stress (SI Appendix Fig. S11). Thus, laser ablation microscopy is a suitable approach to study the microarchitecture of the apical notch region. The same concentrations and incubation times were used for PI staining of gemmae directly on the growth media at 0DaA, 4DaA and 7DaA for SI Appendix Fig. S5.

## Acknowledgments

I would like to thank lab members Susanna Sauret-Güeto, Jenna Rever, Harriet Kempson, Sze Wai Tse and Eftychios Frangedakis for their advice and support. I especially thank Marius Rebmann for providing plant lines and valuable intellectual input. Mihails Delmans developed concepts related to these experiments. Jim Haseloff provided funding and influenced the development of this project. I thank Alexander Jones (Sainsbury Laboratory, Cambridge, UK) for constructive comments on the manuscript, Verena Kriechbaumer (Oxford Brookes University, Oxford, UK) and Monica Perri (University of Oxford, UK) for providing experimental support, Satoshi Naramoto (Hokkaido University, Japan) for providing the 5’ promoter sequence for Mp*YUC2* to create the vector construct, and Kuan-Ju Lu (National Chung Hsing University, Taiwan) for providing data on Marchantia short PIN expression. The Wellcome Centre for Human Genetics Cellular Imaging Core assisted with laser ablation experiments. This work was funded as part of the Biotechnology and Biological Sciences Research Council/ Engineering and Physical Sciences Research Council OpenPlant Synthetic Biology Research Centre Grant BB/L014130/1 to Prof. Jim Haseloff and Biotechnology and Biological Sciences Research Council grant BB/F011458/1 for confocal microscopy to Prof. Jim Haseloff.

## Supporting Information Text

### Transgenic plant lines used for each figure. ET numbers (ET239-PXXX) refer to the Marchantia enhancer trap screening project (1)

Fig. 1 D: L2_239-CsA (ET239-P64) F: L2_239-CsA (ET239-P153) Fig. 2 L2_239-CsA (A: ET239-P21 B: ET239-P21 C: ET239-P153) Fig. 3 L2_239-CsA (A: ET239-P21 B: ET239-P21) C: L2_283-CsA Fig. 4 L2_239-CsA (ET239-P125)

Fig. 5 L2_239-CsA (B: ET239-P153 E: ET239-P153 H: ET239-P125) C, F, I: L2_268-CsA J: L2_283-CsA

Fig. S1 L2_239-CsA (A, B: ET239-P21 C,D: ET239-P33 E,F: ET239-P125 G,H: ET239-P153) Fig. S2 L2_239-CsA (ET239-P153)

Fig. S4 L2_239-CsA (A:ET239-P125 B: ET239-P21 C: ET239-P21 D: ET239-P153 E: ET239-P21 F: ET239-P21)

Fig. S5 Wild Type (Cam accession) Fig. S6 L2_239-CsA (ET239-P125)

Fig. S7 L2_239-CsA (B-E: ET239-P125 G-J, L-O: ET239-P153) Fig. S8 L2_239-CsA (ET239-P21)

Fig. S10 L2_239-CsA (ET239-P125)

Fig. S11 A-F: Wild Type (Cam accession) G-I: L2_239-CsA (ET239-P14) J-Q: L2_239-CsA (ET239-P64)

### Microscopy Methods and Settings

Imaging was carried out using a Leica Sp8 Confocal Microscope and LasX v3.5.7.23225, equipped with a hybrid detector and a pulsed white-light laser, with a HC PL APO 10x/0.40 CS2, a HC PL APO 20x/0.75 CS2 dry objective or a HC PL APO CS2 40x/1.30 oil objective. Images were taken at 1024×1024 frame size in photon counting mode with sequential acquisition, using bidirectional scanning at 600Hz and 2x line averaging. Time gating was active to supress autofluorescence. Imaging for SI Appendix Fig. S5 was done using a Zeiss LSM800 upright confocal microscope and ZEN Blue v2.6, with an EC Plan-Neofluar 10x/0.3 dry objective or a Plan-Apochromat 20x/0.8 M27 dry objective. SI Appendix Fig. S5 confocal images were taken at 1024×1024 frame size with a GaAsP-PMT detector with sequential frame acquisition, using bidirectional scanning at scan speed 8 and 2x averaging. Maximum-intensity projections of the images were obtained from z-stack series sliced at intervals between 1μm-12.5μm. Excitation and collection settings for each fluorophore are given in SI Appendix Supplementary Information Table 1.

Laser ablation experiments were performed using a Leica LMD6000 laser dissection microscope system equipped with a solid state 355nm cutting laser and controlled by Leica LMD6 software. Ablation performed under the 10x/0.3 HCX PL FL objective lens used settings 60 power, 45 aperture, 25 speed; ablation performed under the 40x/0.6 HCX PL FL objective lens used 60 power, 35 aperture, 20 speed. For SI Appendix Fig. S5 laser ablation experiments were performed using a Zeiss PALM MicroBeam Laser Capture Microdissection Microscope with 3 z-step cycles of 3µm; ablations performed under the 10x objective used settings 70 power, 90 focus, 10 speed; ablations performed under the 40x objective used 50 power, 25 focus, 10 speed. All gemmae used were taken directly from the gemma cup of the parent thallus (i.e., 0dpg). Intact gemmae were also taken from the same cups as controls to verify regular growth and marker expression. Gemmae were planted on 50mm agar plates and ablation and imaging performed directly on the plates to minimize mechanical disruption to the plants. For entire notch excision a 90µm diameter circle was drawn with its centre point at the notch apex, and all tissue within this circle was destroyed by laser ablation. All other cells or regions ablated are shown in schematics accompanying the relevant figures.

---

**Fig. S1.**
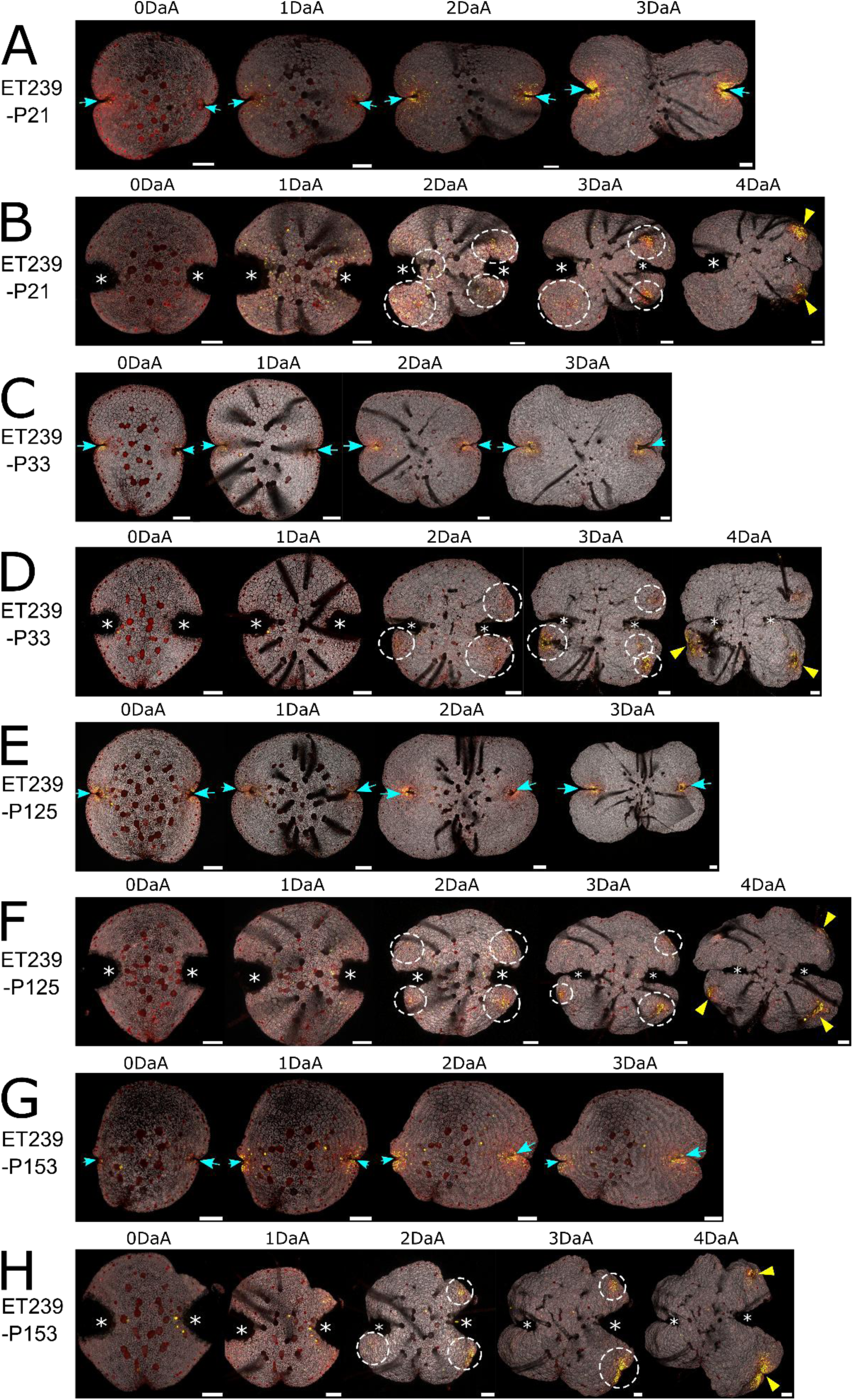
Apical notches and meristem regeneration in the enhancer trap apical notch/meristem marker lines used in this study. (A) is an intact gemma from line ET239-P21 imaged daily from 0dpg until 3dpg. (B) is a gemma from line ET239-P21 with both notches entirely excised by laser ablation, imaged daily from 0DaA until 4DaA. (C) is an intact gemma from line ET239-P33 imaged daily from 0dpg until 3dpg. (D) is a gemma from line ET239-P33 with both notches entirely excised by laser ablation, imaged daily from 0DaA until 4DaA. (E) is an intact gemma from line ET239-P125 imaged daily from 0dpg until 3dpg. (F) is a gemma from line ET239-P125 with both notches entirely excised by laser ablation, imaged daily from 0DaA until 4DaA. (G) is an intact gemma from line ET239-P153 imaged daily from 0dpg until 3dpg. (H) is a gemma from line ET239-P153 with both notches entirely excised by laser ablation, imaged daily from 0DaA until 4DaA. In all lines, mVenus (in yellow) is strongly expressed in the dense cluster of cells in and around the intact and meristematically active apical notch, marked by blue arrows. Reappearance of densely clustered mVenus expression following apical notch excision occurs in all lines (marked by dashed circles) and indicates patches of localized cell division, with newly regenerated notches marked by yellow arrows. The genes whose enhancer elements have been trapped by these lines are unknown (1). All lines express a constitutive mScarlet cell membrane marker (in red) and chlorophyll autofluorescence is shown in grey. The ablation pattern used is as in Fig. 1C. Scale bars= 100μm.

**Fig. S2.**
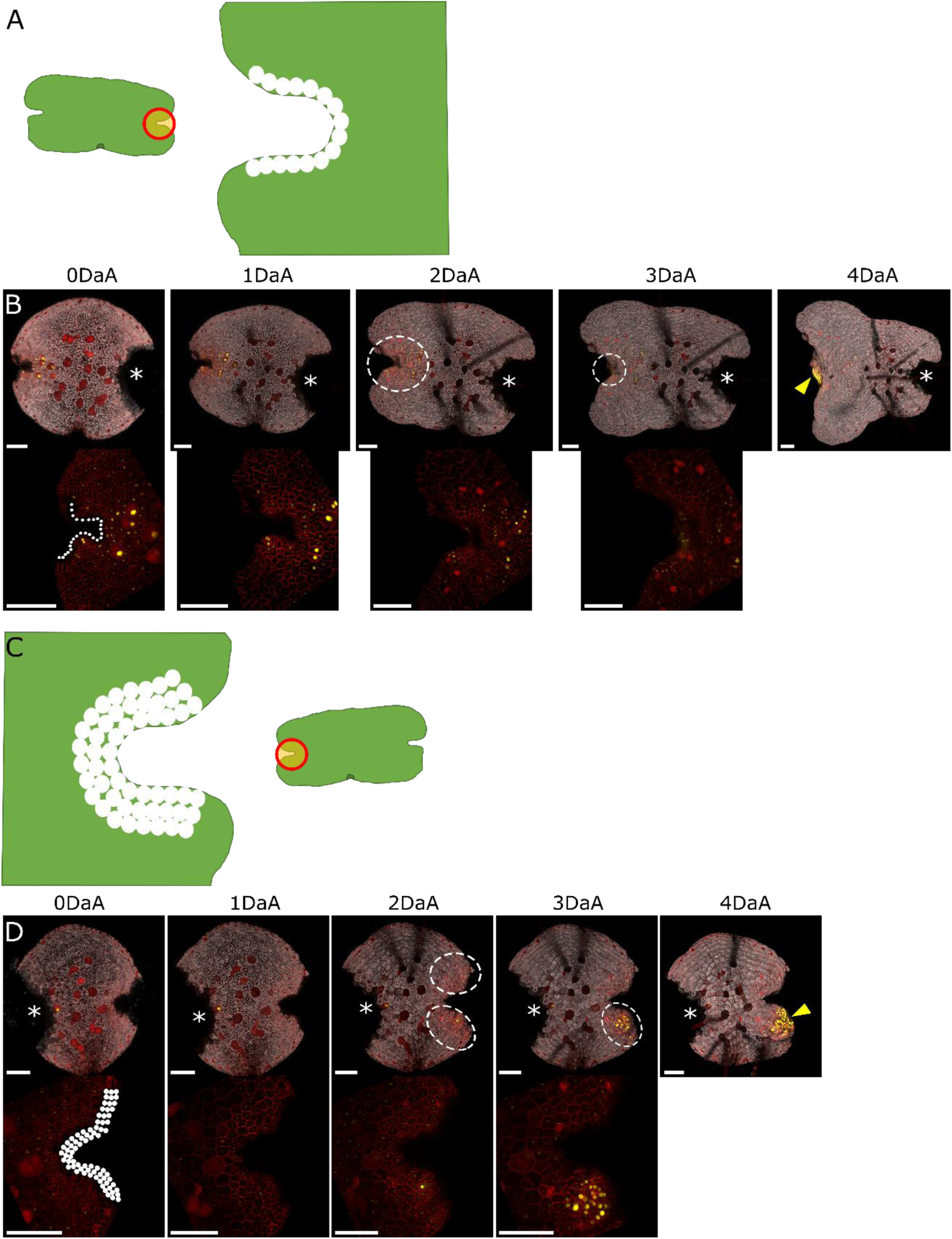
Fine-scale ablation of the apical notch shows that the first three rows of cells are critical in determining meristem regeneration and its location. (A) and (C) are schematics for the ablation patterns used to ablate of the first row and first three rows of cells in the apical notch, respectively. White circles indicate ablated cells, orange bounded by red shows complete tissue excision. (B) and (D) are gemma from the enhancer trap apical notch/meristem marker line ET239-P153 imaged daily during a time course from 0DaA until 3DaA (notch close-up) or 4DaA (whole gemma). White circles mark the position of cells removed by laser ablation in the notch close-up images of 0DaA gemma. One notch in the gemma was entirely excised (marked by asterisk). (B) shows a gemma where only the first row of cells in the notch was ablated. This is sufficient to induce meristem regeneration by 4 DaA, but this occurs at the ablated notch, as seen by mVenus signal marking the localized patch of cell division (dashed circle) and regenerated notch (yellow arrow). (D) shows a gemma where the first three rows of cells in the notch were ablated. This was sufficient to allow patches of localized cell division (dashed circles) and eventually notch regeneration (yellow arrow) to occur elsewhere in the gemma, away from the original notch. Scale bars= 100μm.

**Fig. S3.**
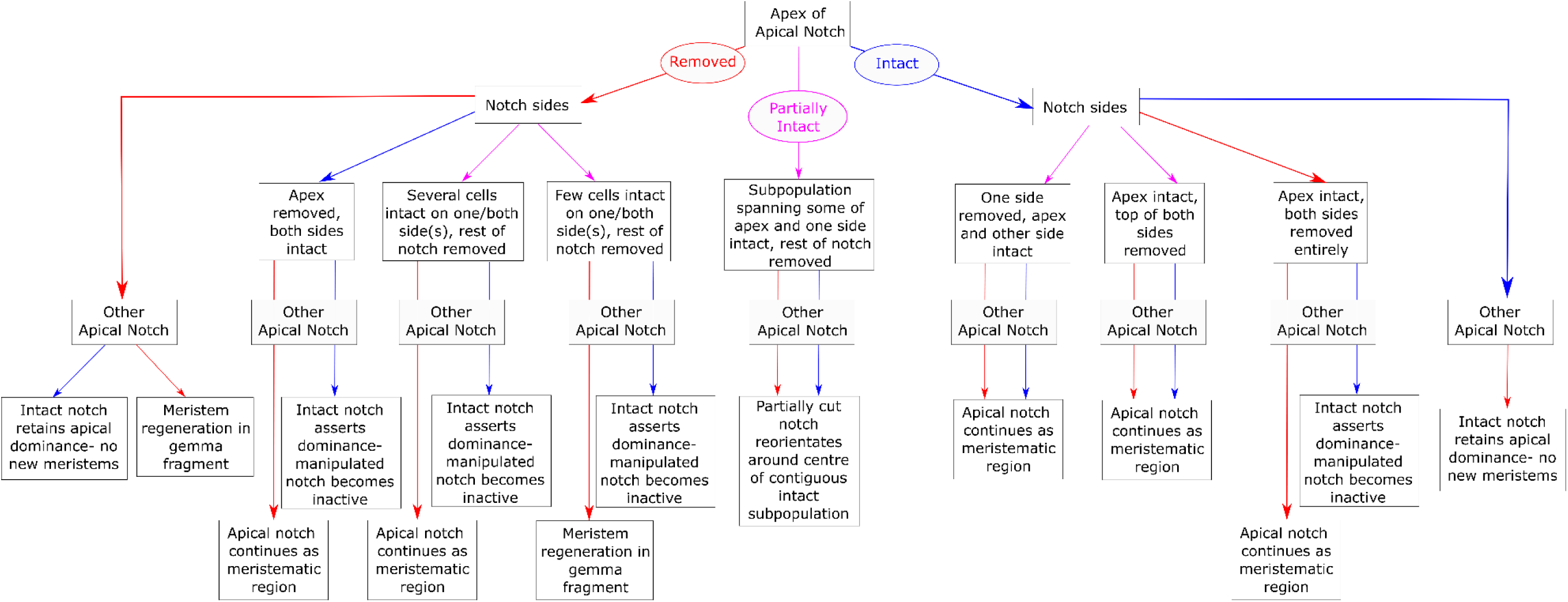
Flow chart of outcomes of ablation experiments. The flow chart progresses from the status of the apex of the apical notch, through the status of notch sides and to the status of the other apical notch in the gemma. Red arrows indicate complete excision, magenta arrows indicate partial ablation, blue arrows indicate where a structure is left intact.

**Fig. S4.**
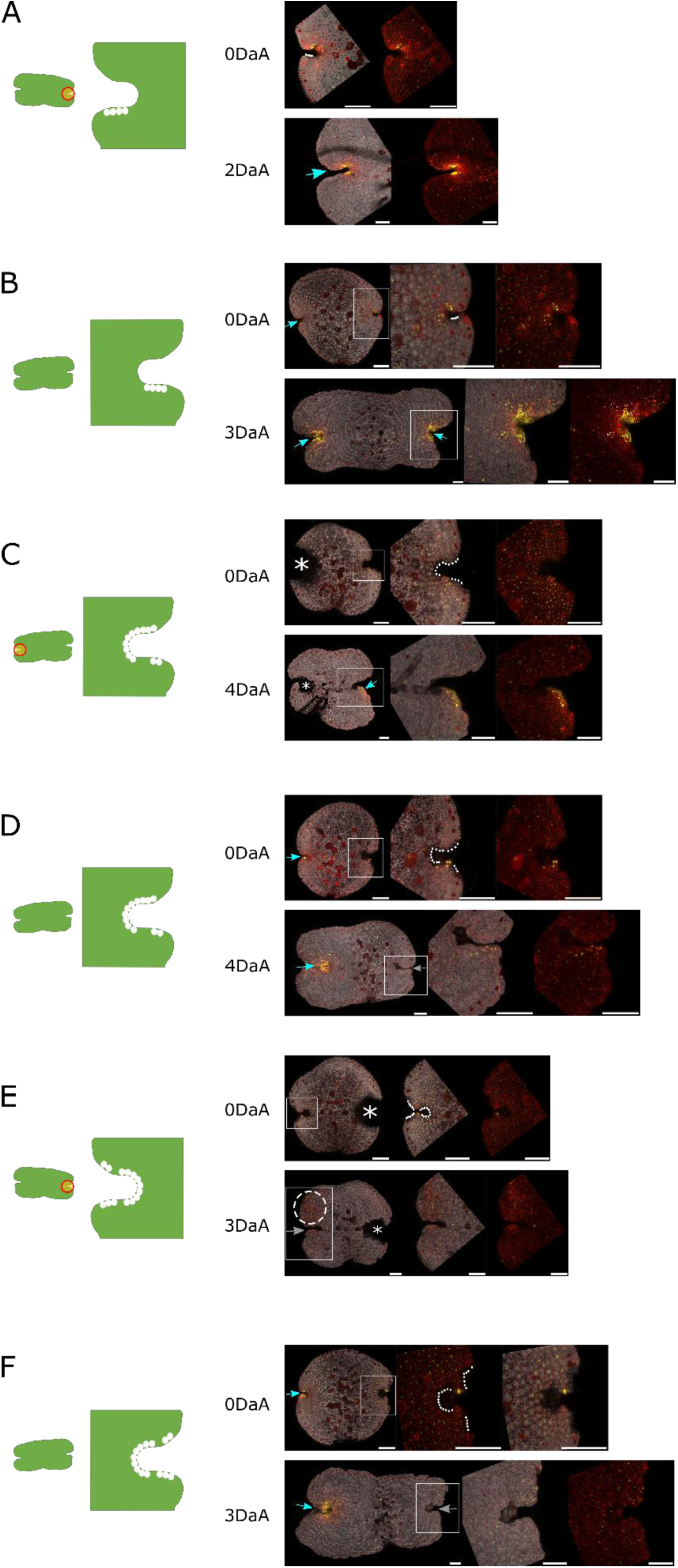
Precise partial ablations of the apical notch. In each sub-figure a schematic of the ablation pattern is given (whole gemma and notch close-up), with white circles marking the location of ablated cells and orange bounded by red denoting excised tissue. Images of the 0DaA gemma are shown on the top row of each sub-figure (whole gemma, close-up of the apical notch area marked in the white box, same close-up without the chlorophyll autofluorescence channel) and the bottom row shows the same gemma at the indicated time point in the same format. Asterisks mark entirely excised notches, blue arrows mark active apical notches, grey arrows mark apical notches that have lost meristematic activity and dashed circles mark patches of localized cell division indicative of meristem regeneration. (A) and (B) demonstrate that ablating one side of the apical notch does not disrupt meristematic function irrespective of whether the other notch in the gemma is intact or not. Ablation of the notch, including the apex, leaving only a portion of one side intact does not disrupt meristem activity if the other notch is excised (C) but if the other notch is intact then meristematic activity ceases (D). Leaving only a few cells intact on both sides of the notch does not sustain meristematic activity, irrespective of whether the other notch is excised (E) or intact (F). If the other notch is excised then meristem regeneration occurs elsewhere in the gemma fragment (dashed circle, E). This is despite the total number of intact stem cells across the whole notch being greater than in the ablation experiment shown in (C). The enhancer trap lines used were ET239-P125 (A), ET239-P21 (B, C, E, F) and ET239-P153 (D). Scale bars= 100μm.

**Fig. S5.**
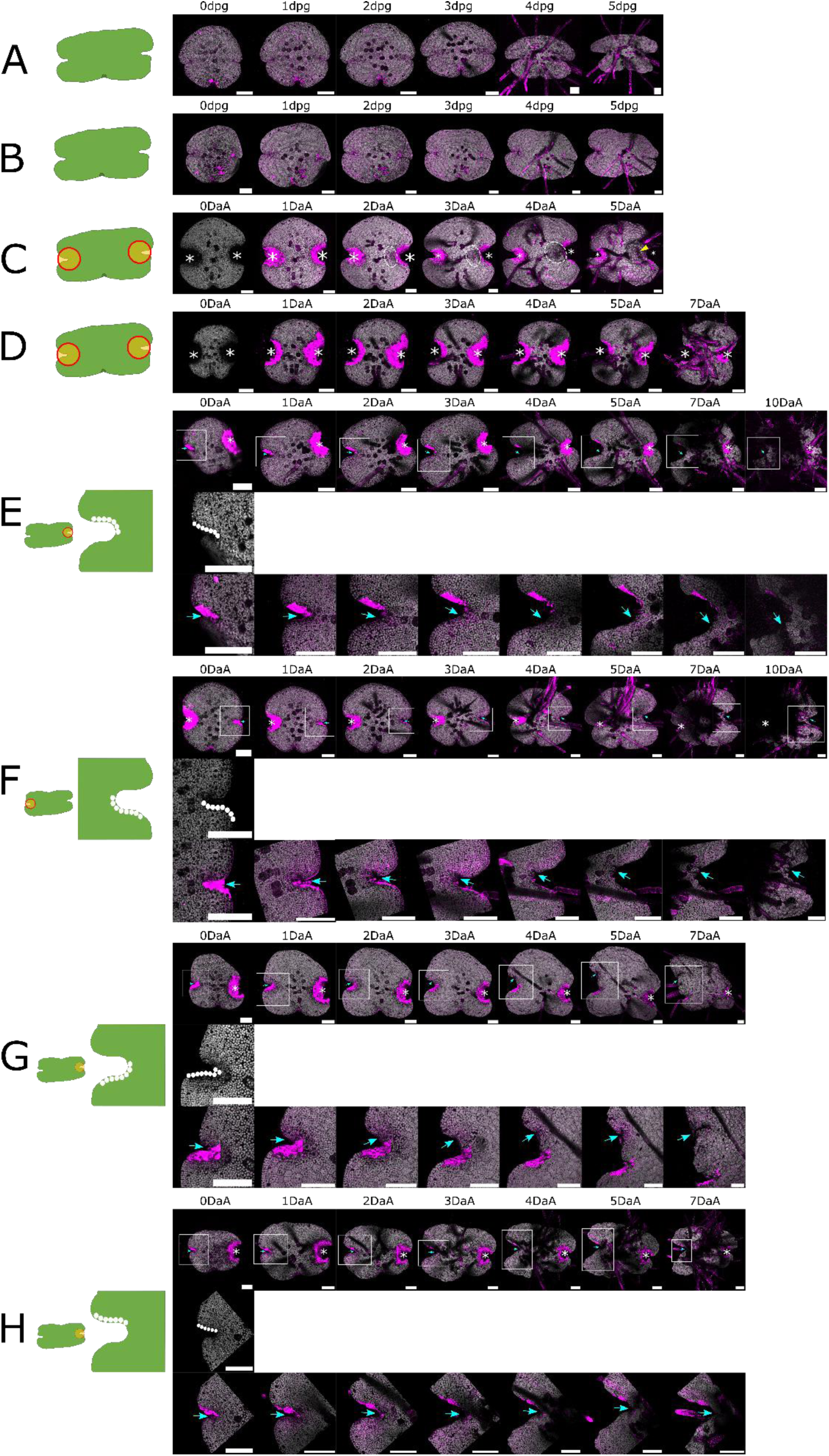
Elevated auxin levels inhibit meristem regeneration but not apical notch reorientation. Gemmae grown on media supplemented with 3µM 1-NAA (A) overall had normal development during the first five days after germination (dpg), compared to gemmae grown on control media (B). The gemmae in (C) and (D) had their apical notches entirely excised (marked by asterisks) by laser ablation according to the schematic shown. While gemma fragments grown on control media (C) showed meristem regeneration (localized cell division=dashed circle, regenerated notch= yellow arrow), no meristem regeneration occurred on gemma fragments grown on media supplemented with 3µM 1-NAA (D). The time courses in (E) and (F) show that elevated auxin treatment did not inhibit apical notch reorientation. The notch changes shape, forming a new apex corresponding to the centre of the original intact subpopulation of stem cells, as in Fig. 3. This occurs at a similar speed to apical notch reorientation in gemmae grown on control media (G). The time course in (H) shows that if the centre of the apical notch is intact then no notch reorientation occurs, indicating that reorientation is not simply because of the generally altered notch morphology observed in gemmae grown in elevated auxin treatments. The schematic shows the ablation pattern used, with white circles marking ablated cells and orange bounded by red denoting excised tissue. Time courses shows the whole gemma, with an asterisk marking the entirely excised apical notch and a blue arrow indicating the position and orientation of the active apical notch. The white square indicates the area shown in the close-ups of the partially ablated apical notch region. All gemmae were wild-type Cam2 accession. Chlorophyll autofluorescence is shown in grey, propidium iodide staining in magenta. Scale bars= 100μm.

**Fig. S6.**
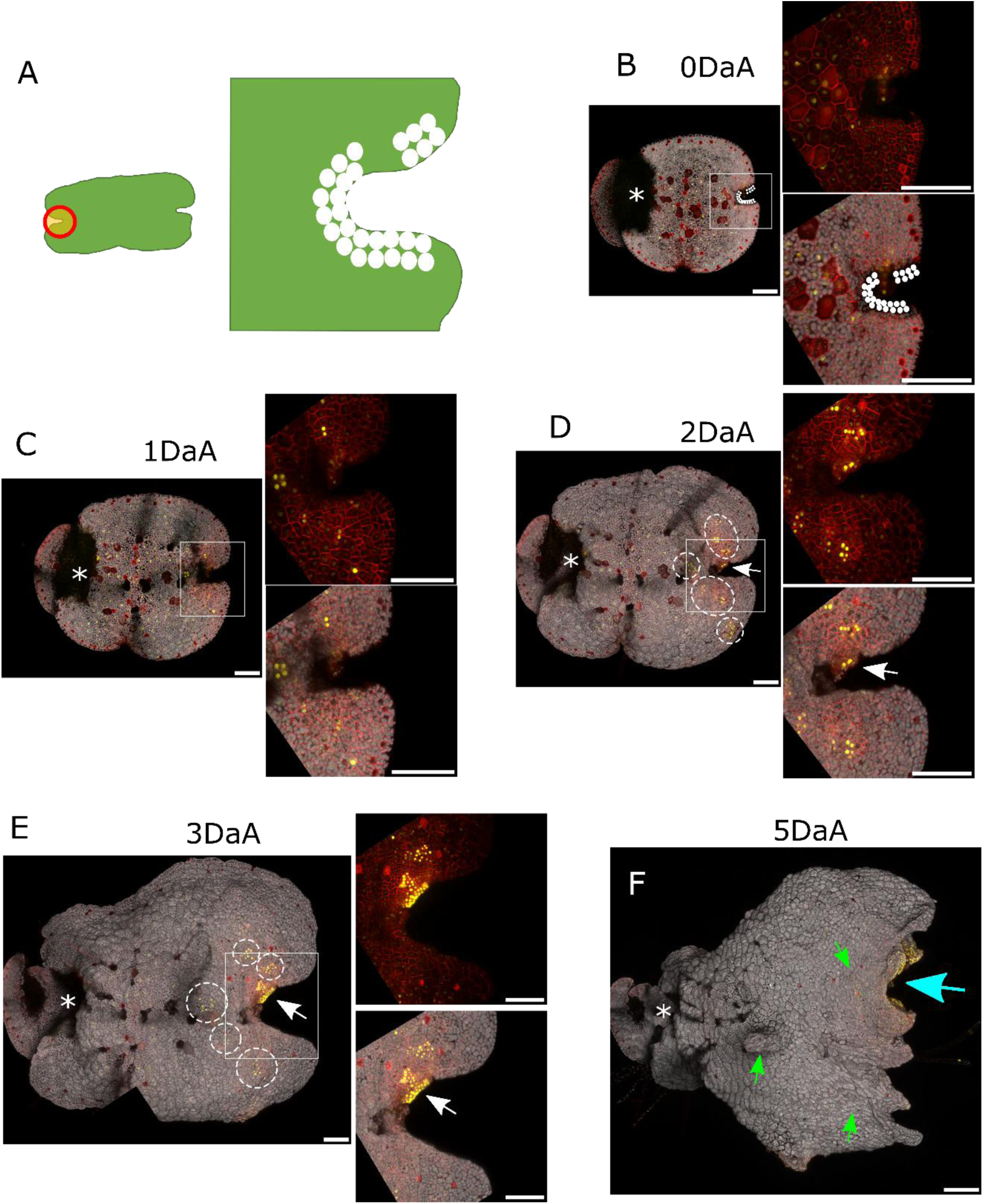
Re-establishment of apical dominance by a partially ablated apical notch. (A) shows a schematic of the ablation pattern used, with white circles marking ablated cells, orange bounded by red denoting excised tissue. (B)-(F) shows a time course of a gemma from the enhancer trap notch/meristem marker line ET239-P125. The images shown are of the whole gemma, together with close-ups of the ablated apical notch region in the white square (shown with and without the chlorophyll autofluorescence channel). Asterisks mark the entirely excised apical notch. (B) is the 0DaA gemma, with white circles marking ablated cells. (C) is the same gemma imaged at 1DaA, (D) at 2DaA, (E) at 3DaA. (F) is the whole gemma imaged at 5DaA. Although the original, partially ablated notch remained active (as indicated by the continuation of mVenus marker signal, white arrows), new patches of cell division emerged at 2DaA (D, dashed circles), with some persisting at 3DaA (E, dashed circles). However, by 5DaA (F) the original notch had reformed and re-established apical dominance (marked by blue arrow), as demonstrated by this notch being the only region of the gemma with mVenus marker signal. The areas that were patches of localized cell division stopped dividing (indicated by loss of mVenus signal) and remained as protrusions on the gemma surface (green arrows). These observations were recorded in one gemma replicate. Scale bars= 100μm.

**Fig. S7.**
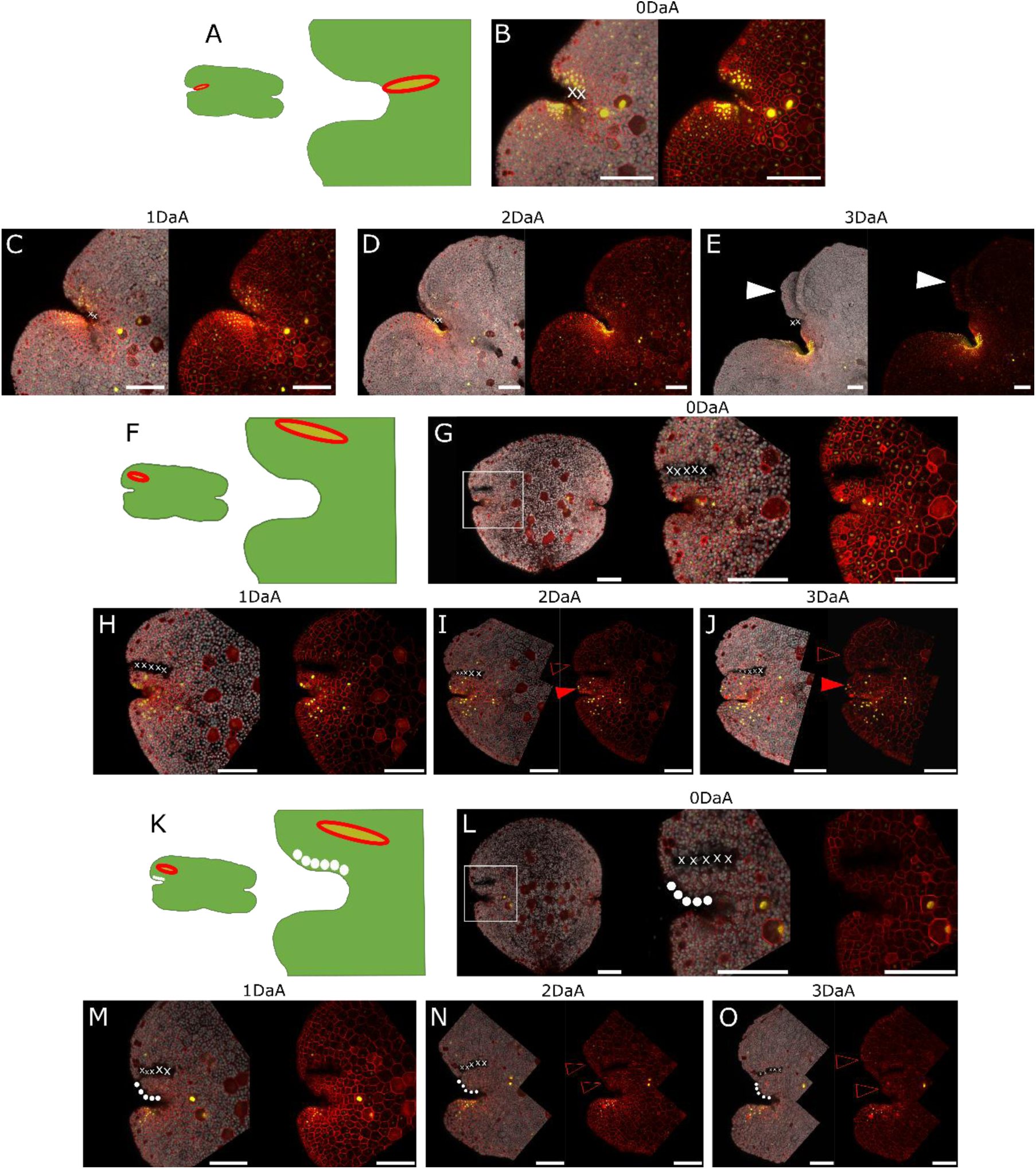
Ablating an incision perpendicular to the apical notch has little effect, whereas ablating an incision parallel to the apical notch alters the balance between cell division and expansion either side of the incision site. (A) Schematic of the perpendicular ablation incision pattern, with orange bounded by red denoting excised tissue. (B) shows a close-up of the ablated notch, with white X marking ablated tissue. (C) is the same gemma imaged at 1DaA, (D) at 2DaA, (E) at 3 DaA. Ablation perpendicular to the apical notch had no obvious effect on notch function, other than the z-axis thallus split (1) in the gemma appearing earlier on the ablated side of the notch (white triangle), at 3DaA. (F) and (K) are schematics of the parallel ablation incision patterns, with orange bounded by red denoting excised tissue and white circles marking ablated cells. (G)-(J) are time courses from 0DaA until 3DaA of a gemma ablated in the patterns shown in (F), with white X symbols marking ablated tissue in (G). Proximal to the incision there was increased cell division (closed red triangle), whereas distal to the incision there was less cell division and more cell expansion (open red triangle), compared to the equivalent position on the non-ablated side of the notch. (L)-(O) is a daily time course from 0DaA until 3DaA of a gemma ablated as in (K), where in addition to the tissue ablated parallel to the apical notch (white X symbols in L) the apical notch cells directly proximal to the incision had also been ablated (white circles in L). In this case the cells on both sides of the incision showed reduced division and increased cell expansion (open red triangles) compared to the equivalent position on the non-ablated side of the apical notch. Gemma images are given in the format whole gemma (G and L only), notch close-up, notch close-up without the chlorophyll autofluorescence channel. Gemmae used were from the enhancer trap apical notch/meristem marker lines ET239-P125 for the perpendicular ablation experiment (B-E) and ET239-P153 for the parallel ablation experiments (G-J and L-O). Scale bars= 100μm.

**Fig. S8.**
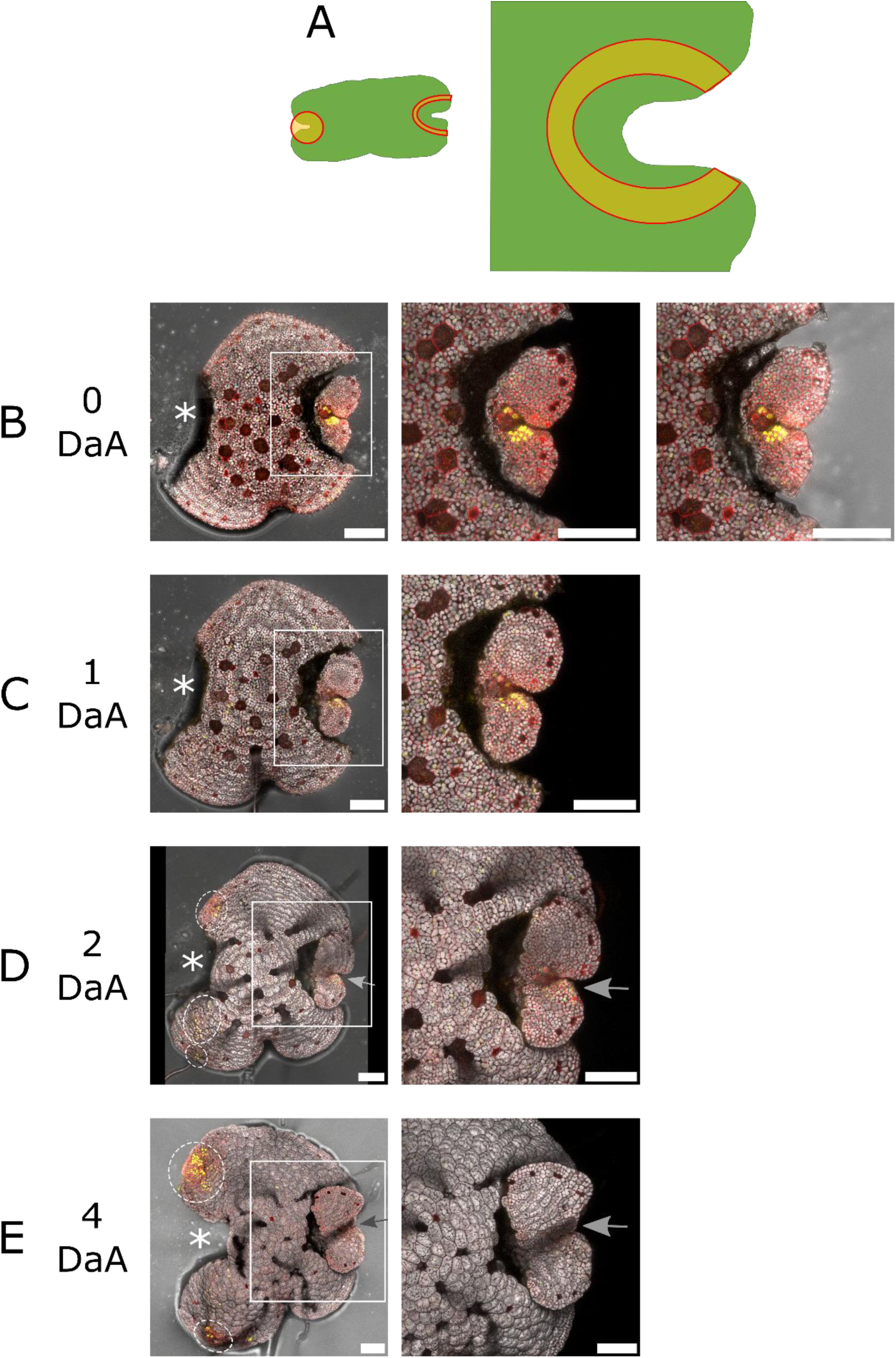
Entirely ablating the tissue surrounding the apical notch results in the isolated notch losing meristematic activity. (A) Schematic of the ablation pattern used, where one notch was entirely excised and laser ablation used to excise all tissue surrounding the other notch (excised tissue shown in orange bounded by red). The isolated apical notch itself was intact and surrounded by a small amount of tissue. The confocal mages show a gemma from the enhancer trap apical notch/meristem marker line ET239-P21 with one apical notch isolated using this ablation pattern, imaged during a time course from 0DaA (B) through 1DaA (C), 2DaA (D) until 4DaA (E). The whole gemma image includes transmitted light PMT channel (gain=315) in addition to the mVenus (yellow), mScarlet (red) and chlorophyll autofluorescence (grey) channels; the white box in the whole gemma image corresponds to the area shown in close-up. The 0DaA close-up includes the transmitted light channel image to confirm that the apical notch was fully isolated from the rest of the gemma fragment by destruction of the intermediate tissue. Appearance of dense regions of new mVenus signal (dashed circles) shows that this isolation removed the effects of apical dominance on the rest of the gemma and allowed patches of localized cell division to emerge. The isolated notch itself showed loss of mVenus signal, cell division ceased and the cells expanded, indicating that this region lost meristematic activity (grey arrows). Scale bars= 100μm.

**Fig. S9.**
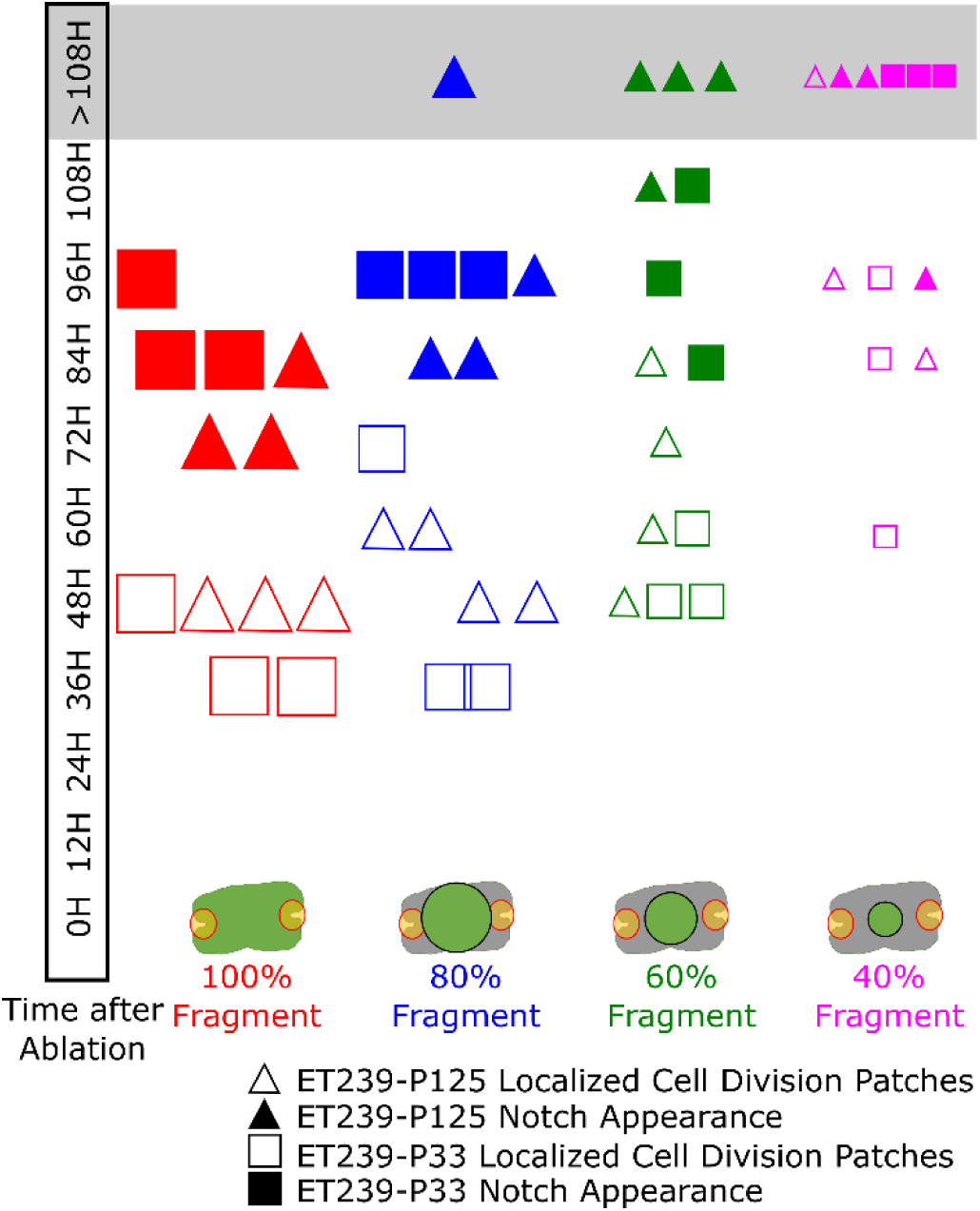
A 108-hour time course using two different enhancer trap apical notch/meristem marker lines shows that larger gemma fragments have faster emergence of patches of localized cell division and faster appearance of morphologically recognizable notches. All 100% gemma fragments showed reappearance of notch morphology within the time course, as did all but one 80% fragment (shown by symbol in the grey >108H box). All 60% fragments had patches of localized cell division and marker signal emerge during the time course, but notches did not appear in all replicates. Only one of the 40% fragments had a notch reappear while one did not even form localized patches of cell division with marker signal. A schematic of the ablation pattern used is given at the bottom, with orange bounded by red indicating an entirely excised apical notch. The black circle shows the laser ablation circle trace used, with the circle diameter calculated as a percentage of the original notch-notch distance. Open symbols denote appearance of patches of localized cell division with enhancer trap marker signal; closed symbols denote when notch morphology within a dense patch of marker signal was first observed.

**Fig. S10.**
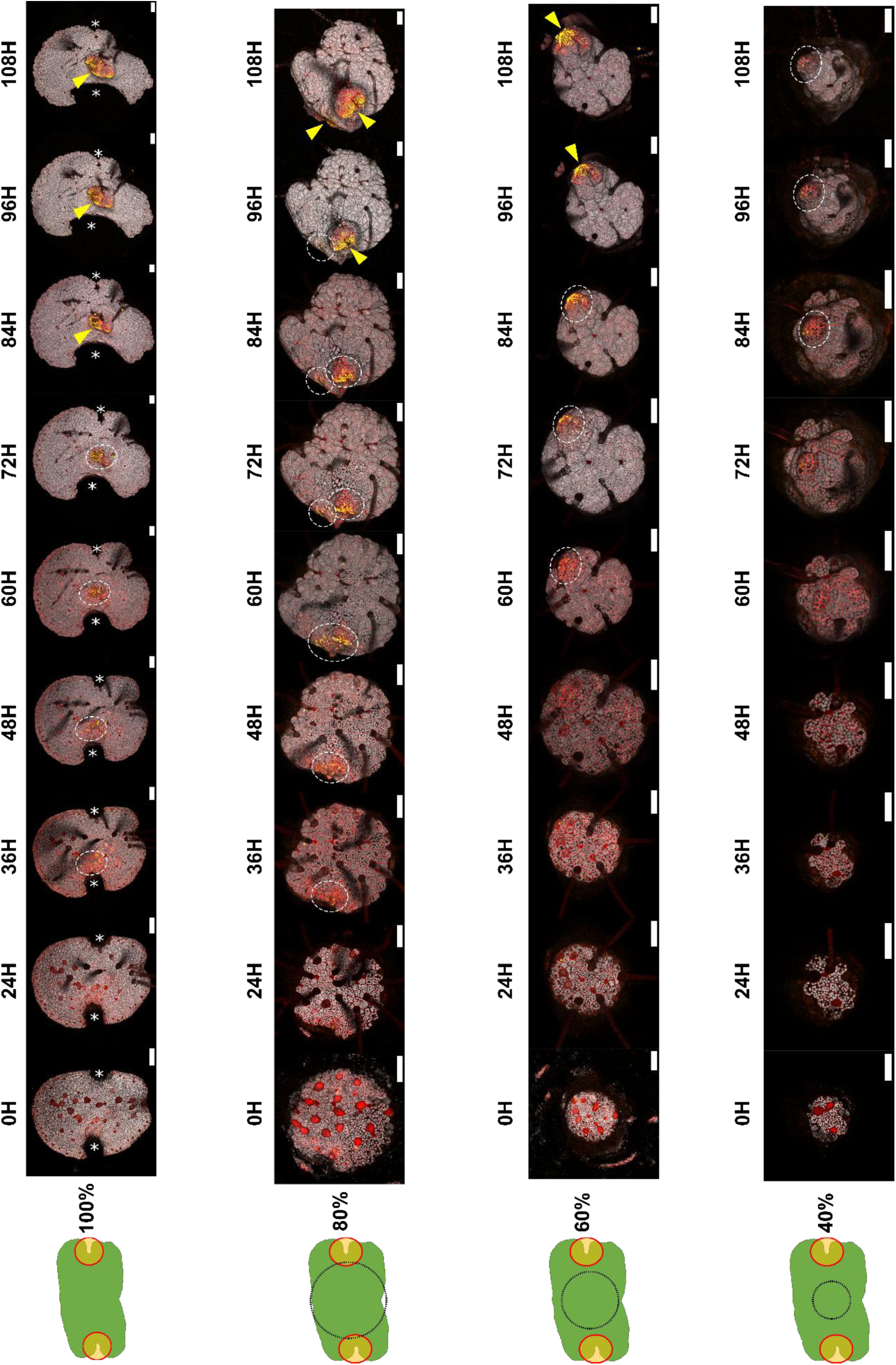
An example time course showing that meristem regeneration proceeds faster in larger gemma fragments. On the far left of each row is a schematic of the ablation pattern used, with orange bounded by red denoting excised tissue in the apical notch removal step (see Methods). The black circle shows the laser ablation circle trace used, with the circle diameter calculated as a percentage of the original notch-notch distance. Only the gemma fragment within the circle trace was retained, with all tissue outside of this destroyed. The emergence of localized patches of cell division, as defined by new, dense regions of mVenus marker signal, is indicated by dashed circles. The appearance of recognizable notch morphology within a region of dense mVenus marker signal is indicated by yellow arrows. These definitions were used to generate the data shown in Fig. 6 and SI Appendix Fig. S9. Both the emergence of localized patches of cell division and notch reappearance occurred faster in larger fragments versus smaller fragments. Gemmae shown were from the enhancer trap apical notch/meristem marker line ET239-P125. Time indicated is in hours after ablation. Scale bars= 100μm.

**Fig. S11.**
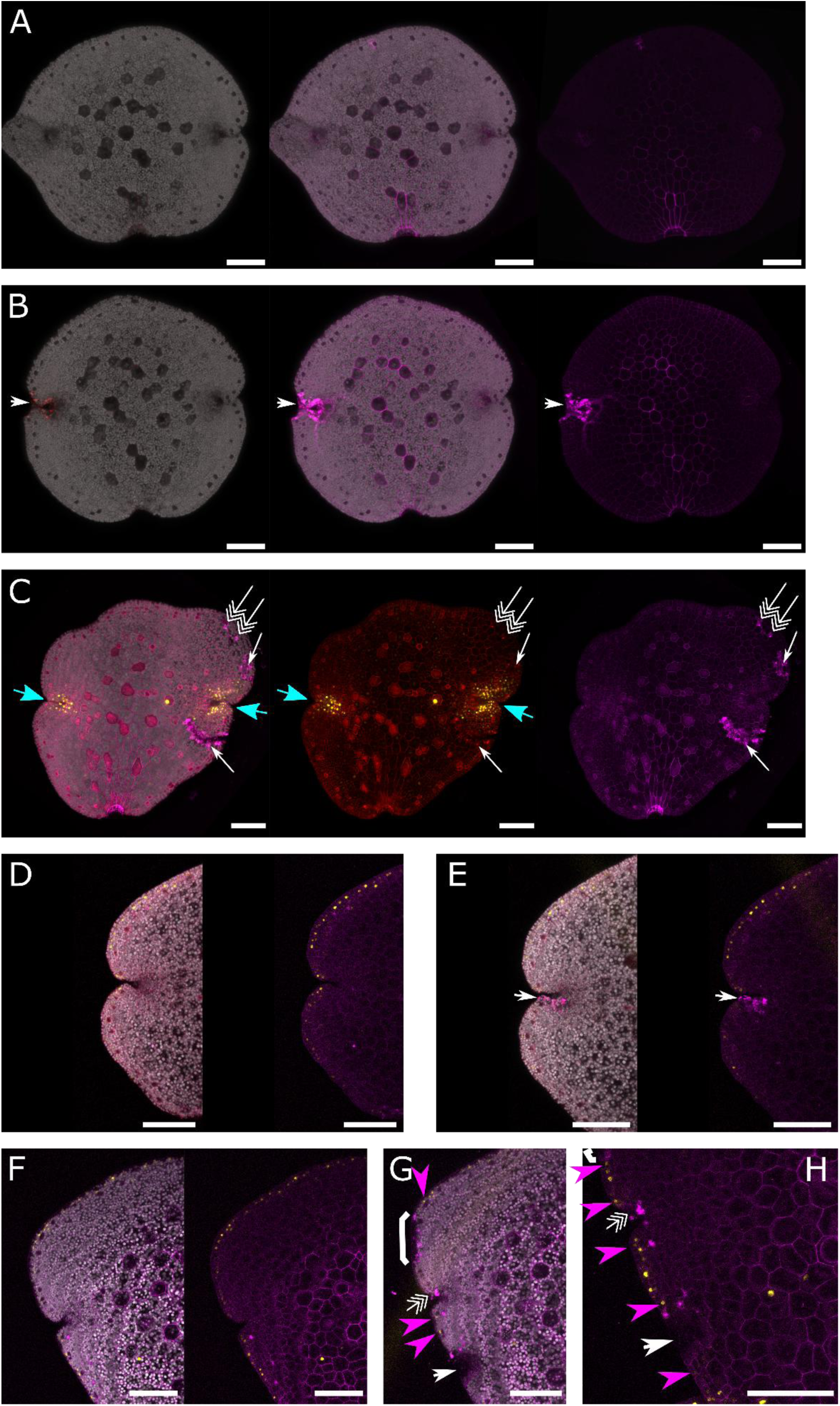
Propidium iodide staining shows that laser ablation kills targeted cells without causing widespread cell death or stress in the gemma. Propidium iodide (PI) staining is shown in magenta. PI stains cell walls but does not penetrate the cell membranes of living cells, however upon cell death it goes into the remains of the cell, staining the nucleus with fluorescence emission in the mScarlet channel wavelengths. (A) shows an intact WT gemma, with no nuclear PI staining. (B) shows a WT gemma whose apical notch was subjected to laser ablation. Targeted cells have nuclear and intracellular PI staining (white arrows), demonstrating that this treatment effectively killed cells in the Marchantia gemma. The damage (also indicated by loss of chlorophyll autofluorescence and mechanical disruption) was restricted to the area targeted by the laser, with neighbouring cells and the wider gemma left intact. (C) shows a gemma from an enhancer trap line (ET239-P14) following various laser ablation targeting treatments. Regions targeted for tissue excision (white arrows) featured multiple other obvious signs of cell damage (loss of chlorophyll autofluorescence, membrane destruction and mechanical disruption) and there were no signs of widespread cell death or stress across the rest of the gemma. Ablation targeting single cells (triple white arrows) only killed those cells. Laser ablation treatment did not have off-target effects on apical notch/meristem mVenus marker signal (blue arrows). (D) shows a gemma region before ablation; (E) shows the same region afterwards. The lower side of the apical notch had been ablated (white arrows in (E)) with no damage caused to the other side of the apical notch. (F) shows a gemma region before ablation, (G) shows the same region after ablation and (H) is a close-up of the same post-ablation gemma. The ablation patterns used were a row of cells at the gemma edge (white brackets), targeting single cells (triple white arrows) and tissue excision (white arrows). Only the targeted areas showed cell death, with agreement between PI staining, loss of chlorophyll autofluorescence and mechanical disruption. Neighbouring cells were unaffected, and enhancer trap marker signal continued unaffected (yellow arrows). (D)-(H) shows gemmae from the enhancer trap line ET239-P64. (A) and (B) show, from left to right, the chlorophyll autofluorescence and mScarlet emssion channels overlaid; chlorophyll autofluorescence and PI emission channels overlaid; PI emission channel only. (C) shows, from left to right, chlorophyll autofluorescence, mScarlet, PI emission and mVenus channels overlaid; mScarlet and mVenus channels overlaid; PI emission channel only. (D)-(G) show, from left to right, chlorophyll autofluorescence, PI emission and mVenus channels overlaid; PI emssion and mVenus channels overlaid. (H) shows PI emssion and mVenus channels overlaid. Scale bars= 100μm.

**Table S1.**
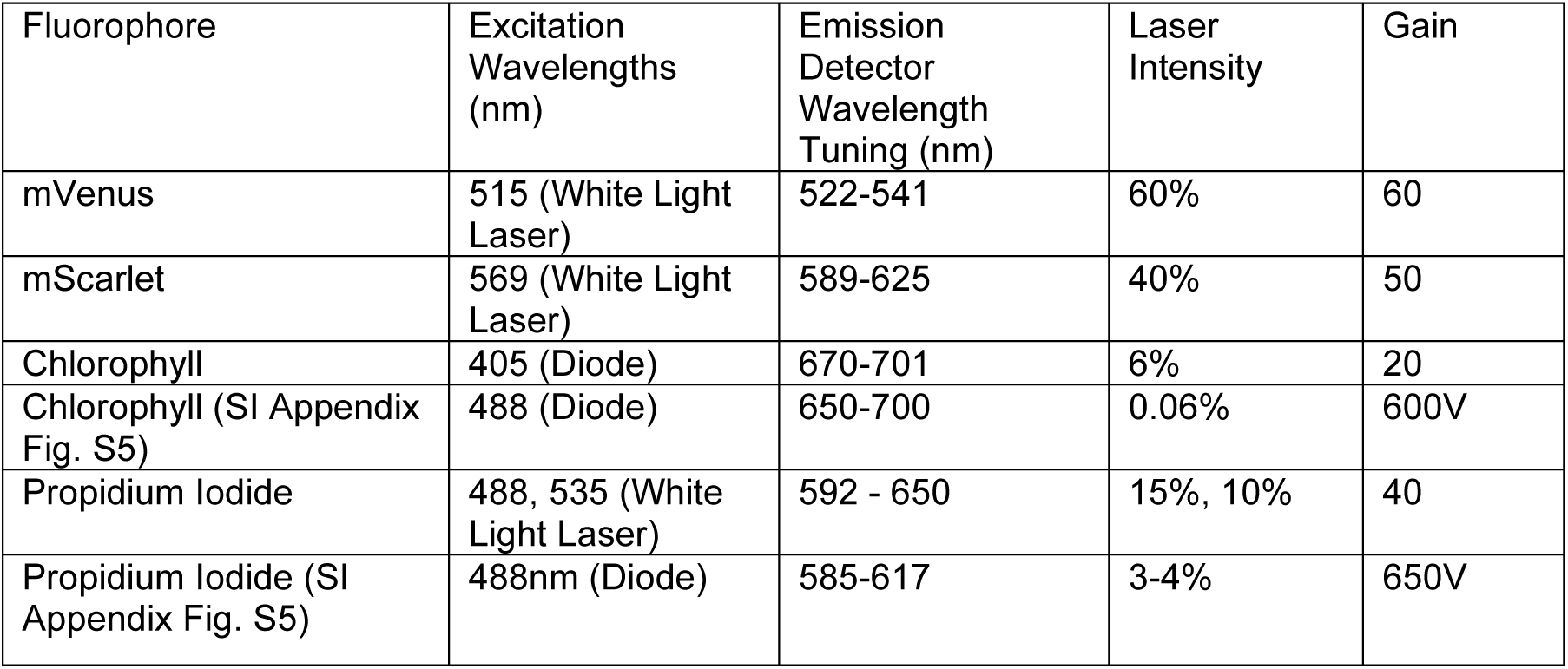
Excitation and collection wavelengths and settings used in confocal microscope imaging.

**Dataset S1 (separate file).** Annotated sequence description of the plasmids used to generate the transgenic lines used in the experiments: L2_239-CsA (16,019 bp), L2_268-CsA (19,174 bp) and L2_283-CsA (26,315bp). The sequences and annotations are provided in Genbank format.

## Notes

### Competing Interest Statement

The authors have declared no competing interest.

### Summary of Updates

SI Appendix Fig. S5 has been updated to include additional experimental data. There are minor edits and clarifications to the Discussion section of the main text.

